# Long range, high-gamma phase coherence in the human brain during overt and covert speech

**DOI:** 10.1101/2021.01.23.427916

**Authors:** Taufik A. Valiante, Bojan Garic

**Affiliations:** Department of Surgery, Division of Neurosurgery, University of Toronto, Toronto, Ontario Canada; Krembil Brain Institute, Toronto, Ontario, Canada; Institute of Biomedical Engineering, University of Toronto, Toronto, Ontario Canada; Electrical and Computer Engineering, University of Toronto, Toronto, Ontario Canada; Center for Advancing Neurotechnological Innovation to Application (CRANIA), University Health Network and University of Toronto, Toronto, Ontario, Canada

## Abstract

Using intracranial electroencephalography (iEEG) in patients undergoing diagnostic workup for epilepsy, we show that during reading, phase coherent high-gamma activity emerges between spatially distant regions of proven involvement in vocalization. Using the novel metric of “phase-dependent power-causal correlations”, causal interactions were shown to be maximal at high-gamma frequencies, and displayed stronger correlations to power fluctuations near the optimal time delay, with this delay potentially accounted for by axonal conduction. We conclude that high-gamma activity may represents propagated, directed, large-scale integration between task related regions of the human brain.

## Introduction

Electrical stimulation mapping (ESM) remains the current gold standard for identifying regions of eloquence in the human brain^1^. ESM has demonstrated particular utility in mapping human language areas, and has revealed multiple critical language regions, where upon ESM, speech arrest, dysarthria, and receptive language problems are observed. An increasingly network oriented approach to understanding brain function^2–6^, suggests that these spatially disparate language areas work cooperatively together in a temporally ordered fashion to generate speech^7^. The signature of such cooperative activity is thought be coherent oscillations^2–4,8,9^, and when such coherent activity occurs between different regions it is assumed to represent a form of communication – so called communication through coherence^3^.

Given that ESM demonstrates necessity of involvement, and coherence demonstrates communication between brain regions, together these lead to a clear and simple hypothesis in the context of language networks – that critical language regions demonstrated by ESM will demonstrate coherent activity during vocalization. As hypothesized we found that the strongest phase coherence could be measured between ESM identified critical vocalization sites in two patient undergoing iEEG recordings. Surprisingly however, the coherence was greatest within the high-gamma range (60-200Hz), and across large scales (centimeters). These findings are particularly unusual since a myriad of data suggests high-gamma activity likely represents the temporal and spatial summation of asynchronous spiking and post-synaptic potentials, and is thus not an ‘oscillation’ that can demonstrate long-range phase coherence^10,11^, although there is recent evidence to the contrary^12^. Despite the discrepancy between our findings and the current understanding that high-gamma is not an oscillation^11^, our data suggests that causal interactions and phase coherence at high-gamma may occur under some circumstances, and play an important role in language networks^13,14^.

## Methods

### Patients

Two patients age 22 and 59 with medically intractable epilepsy underwent craniotomy for implantation of intracranial grid electrodes on their dominant hemispheres. Platinum-iridium grid and strip electrodes (PMT Corp MN, USA) were utilized with 10 mm inter-electrode distances, and 3mm discs. The electrode locations were determined by their pre-operative workups. Stimulation mapping was performed as part of routine pre-resective testing (see **Supplementary Methods** for more clinical details and mapping specifics). Other tasks which included overt and covert reading, and resting with eyes open and closed did not interfere with the clinical ECoG recordings, presented minimal risk to the subjects, and were done as part of a study protocol approved by the UHN Research Ethics Board for which written informed consent was obtained. In both patients, these tests were performed at the completion of their clinical recordings once the epileptogenic, and irritative zones had been determined. Stimulation mapping was performed by applying biphasic current pulses of 100ms duration at 50Hz delivered at various intensities (Ojemann stimulator, Grass Instruments) through various contact pairs. Once an intensity was found to causes after discharges, the current was reduced by 1mA, and applied prior to presentation of an object to be named from a standard set of black and white pictures. The current was terminated when the sentence “This is a {item to be named}” was completed or when a speech disturbance was identified. Stimulation was never applied for > 4 seconds. Currents were further reduced to determine the lowest current required to produce a behavioral response. A site was determined to be necessary if three consecutive errors were induced with stimulation.

### Data acquisition and electrode display

During testing 64 channels of intracranial signals referenced and grounded to a sub-galeal depth electrode were band passed filtered between 0.1 and 1kHz, acquired at 5kHz (Synamps2, Compumedics), and stored off line for later analysis. Recordings were limited to 64 channels and thus for both subjects 32 channels of grid electrodes were sampled with the remainder obtained from strip electrodes in various locations. The acquisition system was of fixed gain in AC mode with a resolution of 3nV/bit.

For 2D display of electrode positions post-operative CT scans were co-registered to pre-operative 3T MRI utilizing Stryker Navigation System. For 3D reconstructions co-registration was performed using the BrainLab system.

### Computational methods

All analyses were performed using MatLab, using custom scripts. None of the electrodes analyzed were part of the irritative, or epileptogenic zones. A common average reference was generated by averaging over all artifact and noise free channels and subtracting this mean from each channel. Noisy channels were identified by visual inspection and power spectra and excluded. For subject 2 analysis was performed only for electrodes within the 2×8 frontal region over which speech alterations were observed. In this subject detailed network analysis could not be performed due to the widespread nature of the irritative zones.

Sixty seconds of contiguous data were used from the four continuous states (**Figure S1**) and included: reading text aloud, reading the same text covertly, rest eyes open, and rest eyes closed. For all subsequent analyses data was decimated to 1kHz after low pass filtering at 500Hz (EEGLab, ‘`eegfilter.m`’). ***For simplicity we refer to subject 1 as an exemplar in the methods***.

To obtain time frequency representations of the neural responses, voltage time series we decomposed using either a complex morlet wavelet transformation (‘`cwt.m`’) using a wavelet number of 5, or band pass filtering the time series and computing its Hilbert transform^15,16^. Both approaches produce a complex valued vector from which instantaneous amplitude and phase were extracted.

The phase coherence (PC) between two signals was computed as previously described^17^:

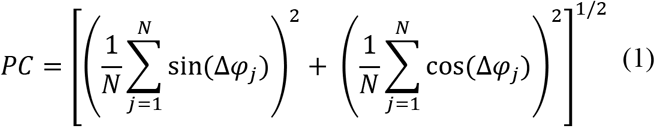

Where Δ*φ_j_* is phase difference between two signals at each of N time points, with N = 60000.

Amplitude envelope correlation (AEC) was computed as the Fischer’s Z transformation of the Pearson correlation between the envelopes of the two time series^18^. Imaginary part of coherence^19^ was computed as follows:

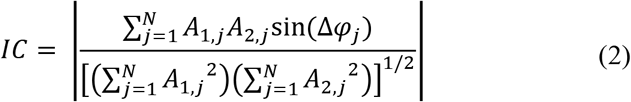

Where *A*_1, *j*_, and *A*_2, *j*_ are the amplitudes of the two time series at each time point. *IC* as originally described^19^ is a signed value, potentially containing directional information. For this measure our interest was to characterize amount and not directionality of phase-lagged synchronization and thus the absolute value was taken.

Frequency dependence of the various coherence measures were computed between 50 and 500Hz in 2.5Hz steps between the unique pairs arising between the face-motor cortex (FMC) and supplementary motor area (SMA) from wavelet transformed time-series. As there were 4 FMC contacts and 3 SMA contacts this amounted to 12 pairs (see **Figure 1**). The frequency dependent coherence profiles were averaged for all pairs, and plotted with their 95% confidence limits obtained by multiplying the standard error at each frequency by 1.96. To compare between conditions, *p*-values were obtained at each frequency utilizing the Mann-Whitney U test (12 samples for each condition at each frequency). Significance was then ascertained from these *p*-values by false discovery rate (FDR) correction^20^. Significant *p*-values fulfilled the following criterion:

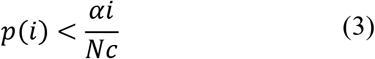

**Figure 1:**
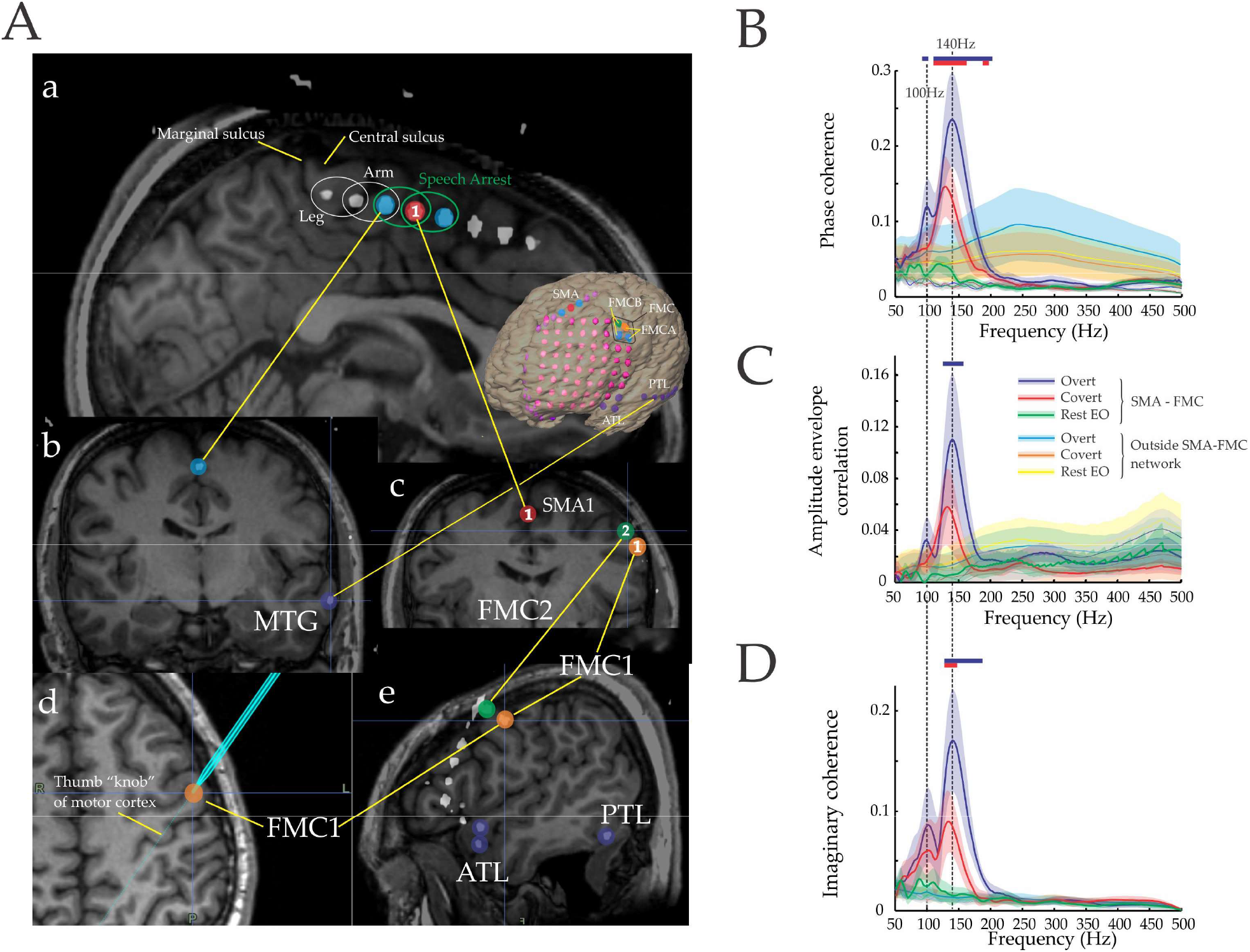
Long-range high-gamma coherence between regions necessary for vocalization. Co-registered pre-operative MRI and post-operative CT, and intra-operative images displaying the location of electrodes within the supplementary motor area (SMA) and face motor cortex (FMC). **Aa-c & e)** Various imaging planes revealing relative spatial locations of electrodes of interest. Those of relevance are four contacts on or near the FMC (collectively referred to as FMC) of which two – FMC1 (orange) and FMC2 (green) are contacts with distinctively high-gamma power increases (FMC1 > FMC2) during reading (**Supplementary Figure S1**); SMA refers to three contacts (red and light blue) in the supplementary motor area, while SMA1 references the single (red) contact which displayed the largest gamma increase of all recorded electrodes during reading (**Supplementary Figure S1**); ATL – three grid contacts on the anterior temporal lobe; PTL – 6 contacts on the posterior aspect of the temporal lobe; and MTG a contact on the middle temporal gyrus. **Ad)** intra-operative navigation referencing FMC1 overlying cortex that is contiguous with the characteristic thumb “knob” of the motor cortex. **B-D)** Frequency dependent synchronization and associated 95% confidence limits averaged over the 12 inter-areal electrode pairs arising from combinations of SMA and FMC electrodes. Synchronization was quantified by computing inter-areal synchronization over a continuous 60s period between the four FMC and three SMA electrodes (a total of 12 unique pairs of contacts) as a function of frequency. Pairs within the two regions (intra-areal) were not included in this analysis (**Supplementary figure 6**). Also shown are average synchronization measures for 200, 12 pair groups selected at random from outside the SMA-FMC network (electrodes ≤ 2cm apart were excluded), as well as the average of time-shifted surrogates of the original (thin colored lines with grey 95% confidence intervals). Two prominent extrema at 100, and 140 Hz are demarcated with horizontal hashed lines. Blue and red bars demarcate regions of significance between overt reading compared to rest eyes open, and covert reading compared to rest eyes open respectively. The precise ranges are tabulated in **Table 1**. Between 2-50Hz only AEC displayed significance and this was noted at 4 and 11Hz at rest.

Where *p*(*i*) are the sorted *p*-values in ascending order and *c* = 1 or a more stringent value of 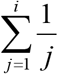 where *i* ∈ *N*, *N* being the number of observations, and α the chosen level of significance (Bonferroni corrected for the number of pair wise comparisons between conditions) set at 0.05. We chose to use the later more stringent condition for tests of significance. Furthermore, for a region to be deemed significant it had to span a range of 5 Hz or more.

To assess statistical significance from chance level coherence, surrogates were generated by rotating the time series by a random number of points (time-shifted surrogates)^21^ then computing the frequency dependent coherence as described above over the same 12 pairs. This method of permuting the time series preserves its statistical nature while obliterating it temporal correlation to its companion time series. Surrogate testing employed 1000 surrogates. Each time a surrogate synchronization was greater than the original time series synchronization a counter was incremented. This was performed across all frequencies. The *p*-value was computed from the equation 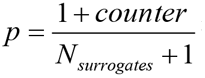^22^ for each frequency. This value was FDR corrected as described above. For subject 1 another type of surrogate was created by randomly selecting a set of 12 pairs from all possible pairs (1891 in total from 62 contacts) 200 times. The same analysis as above was performed although the frequency dependent entrainment for the 200, 12 pair set surrogates were averaged, as was their variances.

#### Phase dependent power correlations

Phase dependent power correlations (PDPC) between channels were computed as previously described^23^. The phase-relation for each frequency was obtained by computing the circular mean 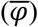 of the phase difference (Δ*φ_j_*) of the two time series and subtracting these quantities 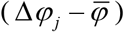. This new time series represented the phase offset on a point by point basis away from the mean phase angle. These phase offsets were binned (along with their corresponding amplitudes from the two time series) into twelve phase intervals of equal size ranging from −π to π, with a central bin spanning from −π/24 to π/24 – the bin that spanned 0 is referred to as has been done previously^23^ as the “good phase”. Within the bins the Spearman rank correlation was computed between the amplitudes of the two signals. Then for each frequency the maximal excursion of a cosine fit was taken as the amplitude of the phase dependent power correlation at that frequency. To determine significance from chance, surrogates were generated by shifting one power time series a random amount of time relative to the other 1000 times, and re-computing the phase-dependent power correlation for each iteration of the random shift.

#### Significance between conditions (‘overt’, ‘cover’, ‘rest’)

To determine significance between different conditions for the various entrainment measures described above, the data was sectioned into 12, 5s “trials”. For each trial the measure of choice was computed yielding 12 samples per condition. *p*-values were obtained using non-parametric statistics (Mann-Whitney U test), and then FDR corrected for multiple frequency comparisons. As the various entrainment measures are not normally distributed a “leave one out” approach was not employed to determine the variance of the sample. The alpha used in the FDR correction was adjusted according to the number of between condition comparisons performed.

#### Power spectra

Power spectra were obtained using multi-taper spectral estimation (‘`pmtm.m`’) over 2s epochs using 2 tapers. Regions of significance between two conditions was assessed in an identical manner to that utilized for frequency dependent synchronization profiles. Furthermore, the same criterion of 5 Hz was used to determine significance within a specified frequency range.

#### Network analysis

To determine the γ_2_ reading network, signals from all channels were band pass filtered between 125 and 165 Hz using a 1000 order windowed FIR filter. The ‘`filtfilt.m`’ function was utilized to avoid phase distortions. Following Hilbert transformation of the filtered signals the instantaneous phase and amplitude were obtained (‘angle’, and ‘abs’ functions respectively). IC was computed for all potential electrode pairs. Statistical significance was determined using bootstrap technique described above using 1000 time shifted surrogates for each electrode pair. The resultant *p*-value was FDR corrected. All possible connection were displayed in matrix form, or reduced to a schematic by collecting only those electrodes that displayed significant synchronization to any contact within the SMA-FMC network.

#### Causal interactions

To assess for causal interaction between different brain regions the adaptive directed transfer function (aDTF)^24^ was computed using the implementation in eConnectome^25^, with bootstrap statistical testing performed by shifting one time series relative to the other and recomputing the aDTF 1000 times. A model order of 12 was determined using the Aikake Information Criterion. The results were qualitatively confirmed using Granger causality analysis^26^. Prior to analyses all signals were demeaned, detrended, and their variance normalized to 1. For subject 1 the DTF values were normalized by the largest value, and then plotted. Normalization was not performed (for visualization only) for subject 2 as the low frequency values for the local channel pair within Broca’s area was considerably larger than for the high-gamma range. The normalization only affected visualization.

#### Phase-dependent power-causal correlations

Phase-dependent power-causal correlations were obtained by first computing time varying causal interactions using the aDTF algorithm in eConnectome^25^ for a contiguous 60s time period, using the same model order as above. To then associate causal interactions, temporal synchrony, and power fluctuations into a single metric a method similar to that of computing phase-dependent power correlations was developed. However instead of correlating the amplitude of the two electrode’s power time series, the source electrode’s power time series was correlated to its outflow, as the aDTF calculation yielded a time series of identical length to that of the original time series. Such an analysis yielded two phase-dependent power-causal plots one for each direction of flow. Bootstrap statistics were performed as described for the phase-dependent power correlations, by shifting the power timeseries a random amount of time relative to the aDTF time series 1000 times and re-computing the phase-dependent power-causal calculation. These statistical analyses were performed for the “good phase”.

#### Cross-frequency coupling

Cross-frequency coupling was assessed by using the phase of low frequency oscillations (2-30Hz in 1 Hz steps) to index the amplitude of the high-frequency (30 to 200Hz by 2 Hz steps) oscillations and measuring the strength of this relationship^27^. To accomplish this the phase interval between −π to π was at which modulation was maximum. The modulation index was computed from this distribution as the Kullback-Leibler distance from a uniform distribution normalized by log_10_^27^. For statistical analyses 200 time-shifted surrogates were generated for each phase-amplitude frequency pair and the modulation index computed for these surrogates. All the surrogate modulation indices were pooled and the surrogate sample divided into twelve equal bins. Within these bins the phases of the low frequency oscillation were used to bin the associated high frequency amplitudes. The entire 60s time series was used. This phase-amplitude distribution was fit to a von-Mises distribution, the mean of which was taken to be the phase angle mean and variance obtained. *p*-values were computed from the normal cumulative density function using the surrogate mean and variance, with *p* values < 0.01 being considered significant^28^. An alternate measure of cross-frequency coupling was computed with the code provided in the supplementary information of Canolty et al. 2006^29^ using a low frequency range of 2 to 30Hz, while setting the high frequency range to 125 to 165Hz. An *mnorm* > 1.96 (p < 0.05) was considered significant.

#### Fits to couple oscillator model

Erickson et al. have described PDPC^23^ as arising from the interaction of two coupled brain regions (oscillators)^30^. They derived the closed form expression of the PDPC curve as a function of coupling weights, and phase delays between the regions for the cases of: 1) bi-directional coupling; 2) unidirectional coupling; and, 3) common drive to both regions. We used these closed form expressions to estimate the coupling strengths between regions as an independent measure of directional influence that may be less sensitive to time delays between time series. For this we performed non-linear fitting of (‘`nlinfit.m`’) the PDPC profiles using the expressions provided by Eriksson et al. for the various coupling regimes^30^.

## Results

We use our first subject as an exemplar, and refer to subject two’s more limited yet confirmatory data set where appropriate. In subject 1 four electrode pairs, representing 7 contacts out of a total of 64 were determined to be critical to speech production (**Figure 1A**). Stimulation of all four pairs (albeit at different current levels) resulted in the subjective feeling of effortful speech, while objectively this was evidenced by speech arrest. Three electrodes were within the SMA, and four within the FMC (**Figure 1A**). The SMA is thought to be a motor planning area, whereas FMC represents the primary output to alpha motor neurons in the brainstem. Both these regions have been shown to be necessary for generating speech^1^. Following mapping the individual was asked to rest quietly with their eyes open and then closed, and then to read overtly and covertly, each for two minutes (**Supplementary figure 1**).

Our primary interest was to determine if long range coherent (synchronous) activity is detectable between regions necessary for the same function. Three measures were chosen for their sensitivities to different synchronization regimes: 1) phase coherence (PC) is an amplitude independent measure of synchrony^17^, 2) amplitude envelope correlation (AEC) correlates the moment to moment amplitude fluctuations independent of the phase^18^, and 3) imaginary part of coherence (IC), which only detects non-zero phase lagged coherence^19^. Clear peaks in coherence were observed within the high-gamma range (**Figure 1B-D**) at 140Hz which was within the γ_2_ range of 125-165Hz (defined in **Supplementary figure 2**), and at 100Hz falling within the γ_1_ frequency range of 96-106Hz. Although the strength of entrainment was variable for individual pairs of electrodes (**Supplementary figure 3**) they invariably displayed a peak within the high-gamma range, similar to the average frequency profile. This was not observed outside the SMA-FMC network (**Figure 1B-D**). These data demonstrate that long range coherent activity emerges between regions engaged in a common task, that this coherence is strongest within the high-gamma range, and occurred at peaks of high-gamma (**Supplementary figures 2 & 4**).

The similar PC and AEC frequency profiles suggest that increases in temporal synchrony (phase) are linked to excitability increases (power). Such correlations between timing and excitability are thought to be the putative mechanism by which effective connectivity between brain regions is regulated through rhythmic gain modulation^3,23^. To explore whether the manifestations of such a mechanism are evident in the human brain, phase-dependent power correlations^23^ were computed for the SMA1-FMC1, SMA1-FMC2 and FMC1-FMC2 pairs of electrodes, representing examples of long range, and relatively local interactions^2^ respectively (**Figure 2A**). During overt and covert reading, increases in phase-dependent power correlation above rest were most strongly observed within the γ_2_ range (**Figure 2B**). In comparison to surrogate time-series, increases in phase-dependent power correlations further extended into γ_1_ frequencies for only the SMA1-FMC1 pair. This was not observed for SMA1-FMC2 the other long range pair. γ_2_ range power correlations were as well observed between FMC1-FMC2, however this short-range pair uniquely displayed correlated activity at rest over a broad range of frequencies (**Figure 2B**). This broad region of correlated activity dissipated during overt reading. Similar robust phase dependent power correlations were observed in our second subject, but in this case between Broca’s area and FMC (**Supplementary figure 4D**). Broca’s area like the SMA is likely a motor planning area^7^. Intermediary sites to Broca’s area displayed weaker correlation to the FMC. Thus effective connectivity at high-gamma frequencies is strongest between functionally related brain regions, and appears not to be limited to short range interactions.

**Figure 2:**
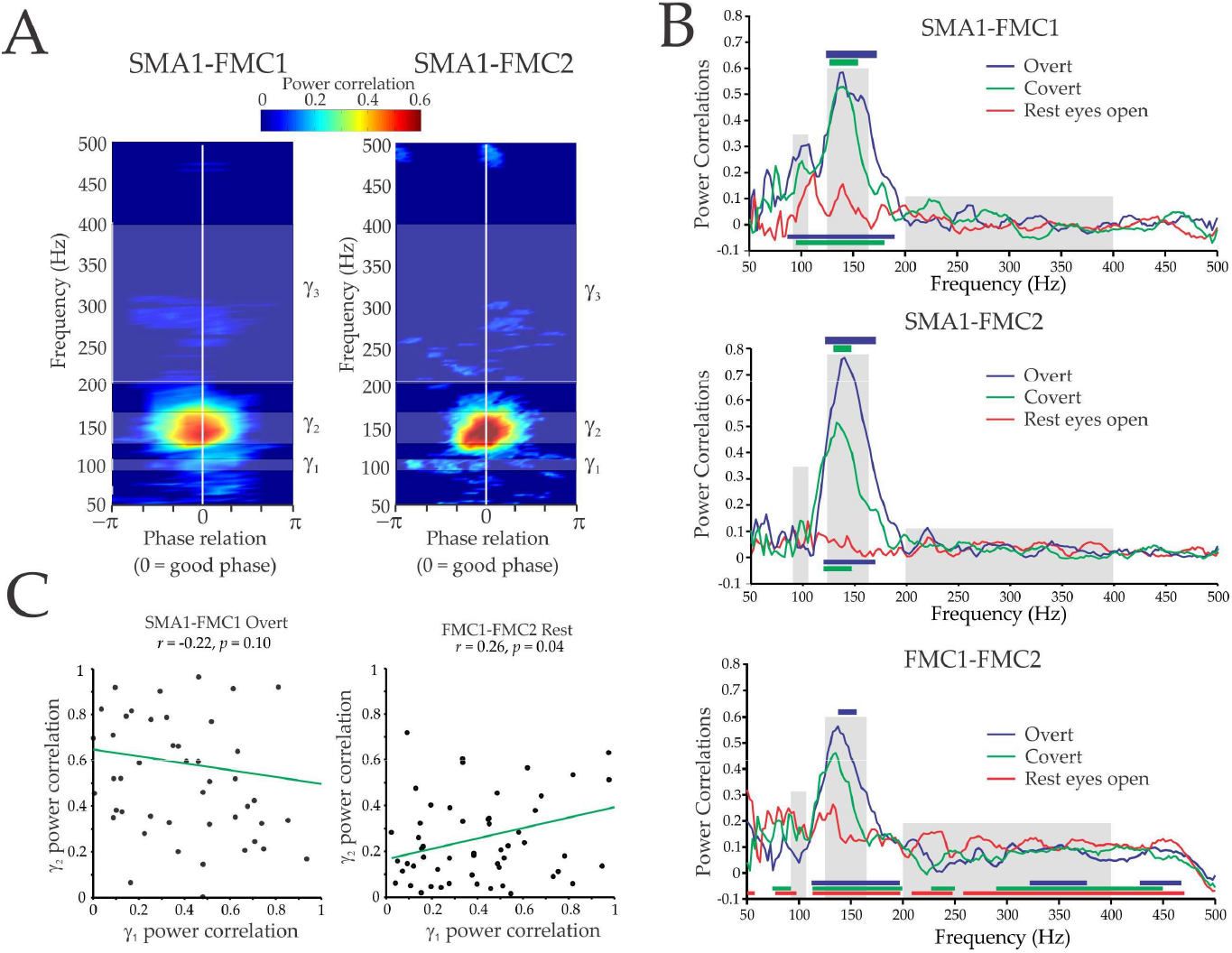
Power and temporal synchrony are comodulated within the γ_2_ frequency range. **A)** Phase-dependant power correlations^32^ between the SMA1, FMC1 and FMC2 electrodes. The various gamma ranges (γ_1_-γ_3_) are shaded in white and labeled on the right. **B)** Plots along the “good phase” relation for various behavioral states. Statistical significance from rest are depicted by the colored bars on top, with their colors corresponding to the state. Similarly the bars at the bottom depict significance from chance. **C)** Power correlations within the γ_2_ range as a function of γ_1_ power-correlations. Green line represents least squares fit. Spearman’s rho (*r*), and *p*-value are shown above the plot. γ_1_ and γ_2_ power-correlations were weakly correlated at rest between FMC1 and FMC2. For all other conditions no correlation was found.

The observed spatial heterogeneity of γ_1_ and γ_2_ phasedependent power correlations suggests that these two high-gamma oscillations may be independently modulated. This was explored by correlating the phase-dependent power correlations of γ_2_ to γ_1_ by segmenting the data into 60 one second epochs. Of the 12 potential pairs in the SMA-FMC network, weak, yet significant correlation was found only between FMC1 and FMC2 in the rest state (*r* = 0.26, *p* = 0.04) (**Figure 2C**). Furthermore task dependent, and frequency specific phase-concentration^2^ (**Figure 3A**) and phase-scattering^2^ (**Figure 3B**) of γ_1_ – γ_3_ were evidenced, suggesting frequency specific communication through coherence and non-communication through non-coherence^3^ can be operative simultaneously. These observations suggest that γ_1_-γ_3_ oscillations may each be generated by different neuronal populations^31,32^ and/or cellular mechanisms. In the human brain awareness and attention can be dissociated by the frequency of gamma oscillations^33^, suggesting that the neural correlates of behavior can be parcellated by frequency^4,33,34^. Our data support such frequency specific segregation, and further suggest that neural activity within the high-gamma range can be indexed by modulations of coherence.

**Figure 3:**
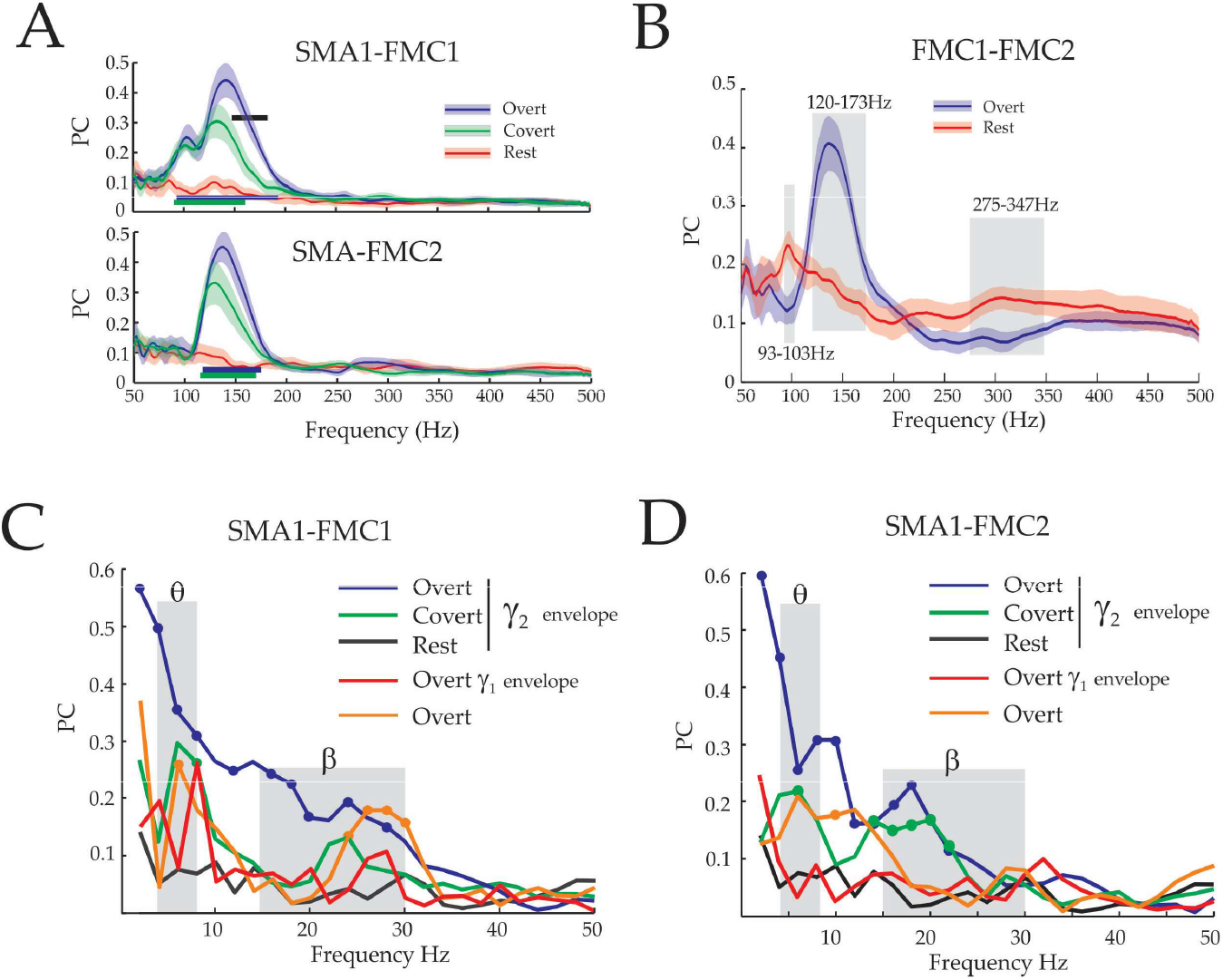
Long range γ_2_ phase synchronization and cross frequency coupling. For A) and B) average PC and associated 95% confidence intervals computed from 12, 5s intervals and then averaged for SMA1-FMC1 and SMA1-FMC2. Significance between overt and covert reading is depicted with a black bar whereas difference as compared to rest are depicted with color bars below, with the color of the bars corresponding to behavioral state. **A)** Between SMA1-FMC1 there is independent modulation of γ_1_ and γ_2_ by task, whereas SMA1-FMC2 only displays phase synchronization within γ_2_ that is not task modulated. **B)** PC between FMC1-FMC2 increases from the rest state within the γ_2_ range during overt reading while simultaneously decreasing with the γ_1_ and γ_3_ ranges. Grayed areas identify regions of significant differences. **C&D)** Frequency dependence of amplitude co-modulations of γ_1_, γ_2_ envelopes and LFPs between SMA1 and FMC1&2. Symbols where present identify significant increase from chance. Grayed areas for C) and D) identify θ and ß ranges.

Coherence within a specific frequency band is but one mechanism of coupling excitability. Cross-frequency coupling^29,35,36^, is another mode of coherence which links excitability between different populations of neurons, or different processes within the same population at different time scales. In the context of human language, such cross-frequency coupling has been suggested to be a means of transforming sensory stimuli into neural syntax^37,38^. To explore this mechanism the PC of the envelope of the γ_2_ oscillations between the SMA1 and FMC1&2 were computed as a function of frequency (2-50Hz). During overt reading γ_2_ power at FMC1&2 was modulated in phase with γ_2_ power in SMA1 at frequencies below 30Hz (**Figure 3C&D**). In stark contrast to this broad pattern of phase coherence, during covert reading γ_2_ envelopes were punctuated at theta^38^ and beta frequencies. Low frequency oscillations themselves only displayed significant phase coherence at theta (4-8Hz), and beta (15-30 Hz) frequencies during overt reading-not as broadband as that observed for the γ_2_ envelopes during overt reading (not shown). Despite the strong low frequency modulations of the γ_2_ envelopes, these modulations were not phase coherent with low-frequency oscillations (**Supplementary figure 5**). This was confirmed as the modulation index^27^, and ‘mnorm’^29^ (computed for the canonical low frequency ranges and the ranges over which significance was observed in **Figure 3C**) were not significant (not shown). Overall while γ_2_ was synchronously modulated at low frequencies between the SMA and FMC, the frequency profile of this modulation was different for overt and covert reading. Such envelope phase coherence in the context of the observed phase-dependent power correlations suggests that phase-relations are not stochastic, but are temporally organized at low frequencies^29,36,37^. This would explain the behaviorally distinct amplitude modulation of high-gamma activity, and suggest the imposition of correlations and decorrelations^39^ at low frequencies by other regulatory brain regions^4^. Such modulations may reflect/determine the behavioral state/fate of high-gamma oscillations: those broadly modulated at vocal formant frequencies (< 50Hz)^40^ being destined for overt speech, while those more narrowly punctuated^37^ destined for covert speech^38^.

**Table 1:** Regions of increased synchronization for inter-areal electrode pairs within the SMA-FMC network. Overt and covert reading states were compared to the rest eyes open state. Behaviorally modulated synchronization regions are discerned within the high-gamma range. No significant **intra-areal** synchronization was detected (**Supplementary figure 6**). Peak frequencies are presented as mean ± SD, and significance between conditions as *p*-values (Mann-Whitney U).

| Synchronization measure | Inter-areal synchronization (n = 12 pairs) |  |
| --- | --- | --- |
|  | Overt > Rest eyes open | Covert > Rest eyes open |
| PC | 93-103Hz<br>110-203Hz<br>Dominant peak: Overt - 140Hz ± 0.76, covert - 130Hz ± 0.6; $p = 6 \times 10^{-5}$ | 110-163Hz<br>188-198Hz |
| AEC | 125-158Hz<br>Dominant peak: Overt - 141Hz ± 1.6, covert - 135 Hz ± 2.4; $p = 0.007$ | N.S |
| IC | 128-193Hz<br>Dominant peak: Overt - 140Hz ± 2.4, covert - 137 Hz ± 1.96; $p = 0.2$ | 128-148Hz |

The few regions considered in the previous analyses represent a small fraction of a larger fronto-temporal language network that the SMA and FMC are known to be part of^7^. To determine if we could extract such a network, IC was utilized to capture the characteristic phase and power correlations that distinguish the interactions within the SMA-FMC network (**Figure 1D**). IC as well provided the greatest “contrast” between electrode pairs within the SMA-FMC network, and those outside the network (**Figure 1D**). IC was thus computed between all possible pairs of electrodes within the γ_2_ range. The results of this analysis (**Supplementary figure 6**) were summarized by reducing all possible significant combinations to a schematic – of those connected directly to the *a priori* SMA-FMC network and those that were not (**Figure 4**). During overt speech production (**Figure 4** & **Supplementary figure 6**) frontal and temporal regions, regions of the brain known to be involved in word production/syllabification (pre-central and inferior frontal gyri), and self-monitoring (superior temporal gyrus) of overt speech respectively^7^ cohered with the SMA-FMC network. During covert speech many of the same regions remained coherent with the SMA-FMC network, with the intersection of these two graphs revealing a “core” network of pre-frontal areas^7^, and rather conspicuously a single middle temporal gyrus contact (MTG – **Figure 1B**, **Figure 4A**) – a region of the brain involved in lemmal retrieval and selection of the lexical phonological output code^7^. This contact did not display significant power increases (**Supplementary Figure 2B**) within the γ_2_ range despite its coherence to the SMA-FMC network (**Figure 4B, S1 and S7**). A similar network was obtained for γ_1_ range activity during overt and covert reading (**Supplementary Figure 8**). Interestingly sub-network analysis revealed non-zero phase lag coherent activity between the frontal (excluding the SMA-FMC network) and temporal lobes exclusively within the γ_1_ range, while simultaneously being phase-coherent to the SMA-FMC network within the γ_2_ range (**Supplementary Figure 7**). The similarity of the γ_2_ network and known language networks^7^ suggests γ_2_ is a reliable marker of language network formation. Furthermore these data suggest that frequency specific and spatially heterogeneous “transmission lines” and multiplexing can occur within transient human brain networks^41^.

**Figure 4:**
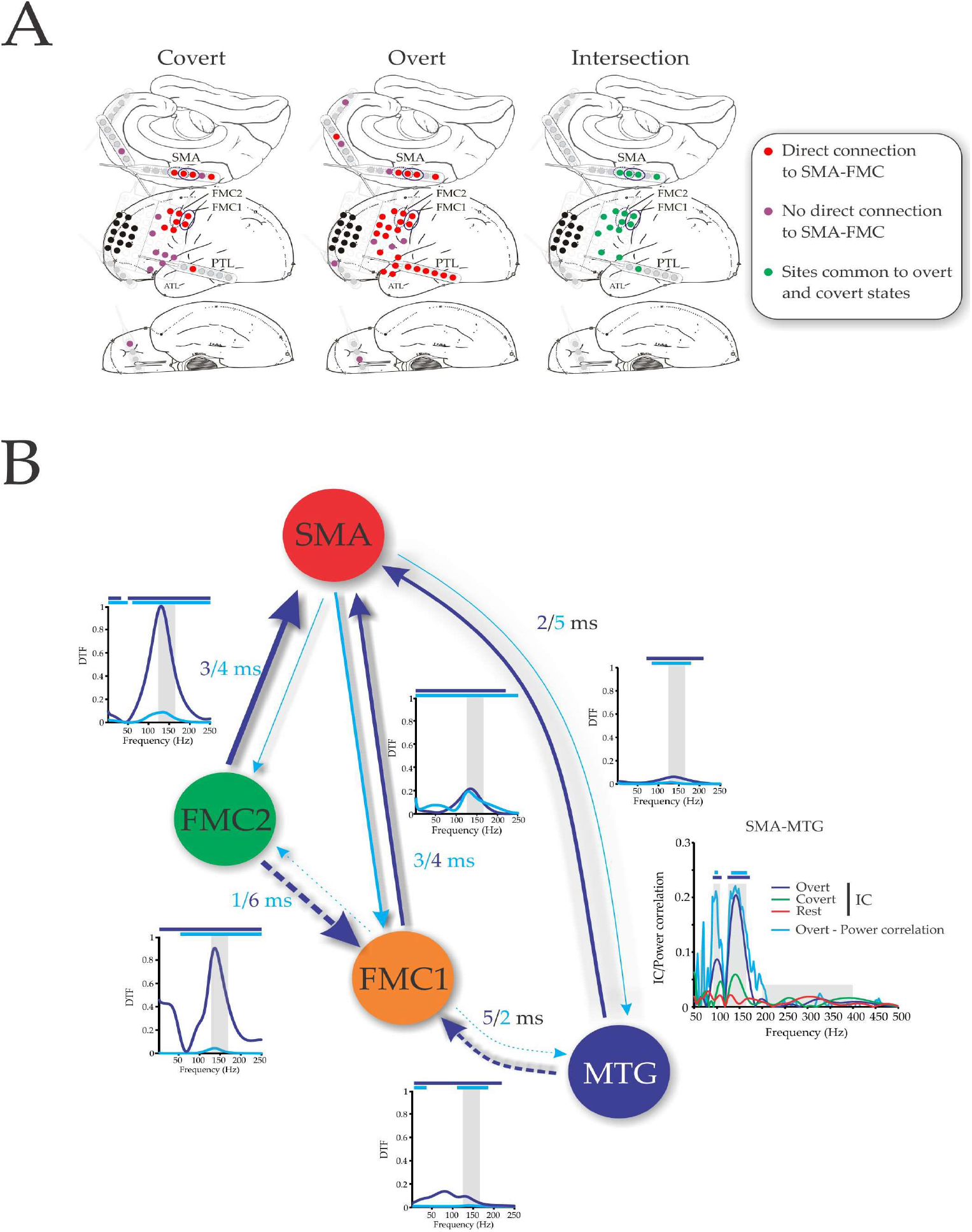
A γ_2_ reading network. **A)** IC was computed between all possible contact pairs. Electrodes entrained directly (red electrodes) or indirectly (purple electrodes) to the SMA-FMC network are shown during overt and covert reading, and the intersection of these two networks (green electrodes). **B)** Four distinct contacts in three distinct brain regions were analyzed for their time delays and causal relationships. MTG was selected from the analysis in A). As well is shown the frequency dependant IC and phase-dependant power correlations profiles between SMA and MTG. Significant regions are represented by horizontal bars above the plots and correspond in color to the legend. The time delays were obtained from the circular mean of the good phase within the γ_2_ range, and their colors correspond to the arrow colors. Forward and backward directions sum to 7ms, the time for a full cycle at 145Hz. The magnitude and direction of the directed transfer function (DTF) between sites is coded in their respective arrow’s directions and color. Bars above depict frequency ranges of significance. The dotted arrows represent questionable “direct” interactions. Grayed bars corresponds to the γ_2_ frequency ranges.

The identification of such a widespread non-zero phase lagged interactions implies time delayed transmission between network nodes. It is known that during object naming different regions of the network identified above are engaged in a temporally ordered fashion^7^. To determine if the time delays within the γ_2_ range correlate to what is known about the temporal organization of this network^7^, as well as to known conduction times between brain regions, we focused on mono-synaptically connected nodes of the reading network. Monosynaptic and reciprocal connections have been identified between SMA and FMC^42^, and very recently monosynaptic connections between regions homologous to the middle temporal gyrus (superior temporal sulcus) and SMA (6DR/C)^43^ have been described in non-human primates. We used the circular mean of the phase difference between contact pairs to estimate the time delay between mono-synaptically connected regions. This in fact is the “good phase” used in the phase-dependent power correlation analyses^23^. The time delays between SMA and FMC were under 6 ms, consistent with the estimated upper limit for conduction time between SMA to FMC in humans^44^. However, whether these delays represent a lead or lag cannot be easily disambiguated. In an attempt to constrain the directedness of these delays, the source of maximal causal interaction between two regions as determined by the aDTF was computed (**Figure 4B**) and qualitatively confirmed using Granger causal methods (not shown). Consistent with its early involvement in language processing^7^ MTG outflow always exceeded inflow (**Figure 4B**), constraining MTG to leading SMA1 and FMC1 in phase. The symmetrical causal interactions between FMC1&2 and SMA1 was consistent with known bidirectional anatomical^42^ and functional^44^ connections between these regions. However since causal interactions were greater for FMC1→ SMA1, FMC1 was taken to lead SMA1. Rather conspicuously the apparent causal interactions within this network were maximal within the high-gamma range (**Figure 4B** – also see **Supplementary figure 4 for subject 2**).

Since measures of causal interaction can be confounded by time delays^45^ we sought alternative confirmatory evidence for the bi-directionality of the interactions that between FMC and SMA, and additionally to provide collateral support to the suggestion that causal outflow was greater from FMC towards SMA than vice-versa (**Figure 5**). Based on the assumption that phase-dependent power correlation arise from the linear interaction of two coupled oscillators, closed form solutions have been derived for the phase-dependent power correlation dependent on unidirectional, or bidirectional interactions^30^. On the whole the birectional model provided qualitatively better fits than the unidirectional model (**Supplementary Figure 9**) except for the relationship between MTG and SMA (**Figure 5**). These independent estimation of directional influence, support bidirectional interactions within the FMC-SMA network and support a great directional influence from FMC to SMA than the reverse.

**Figure 5:**
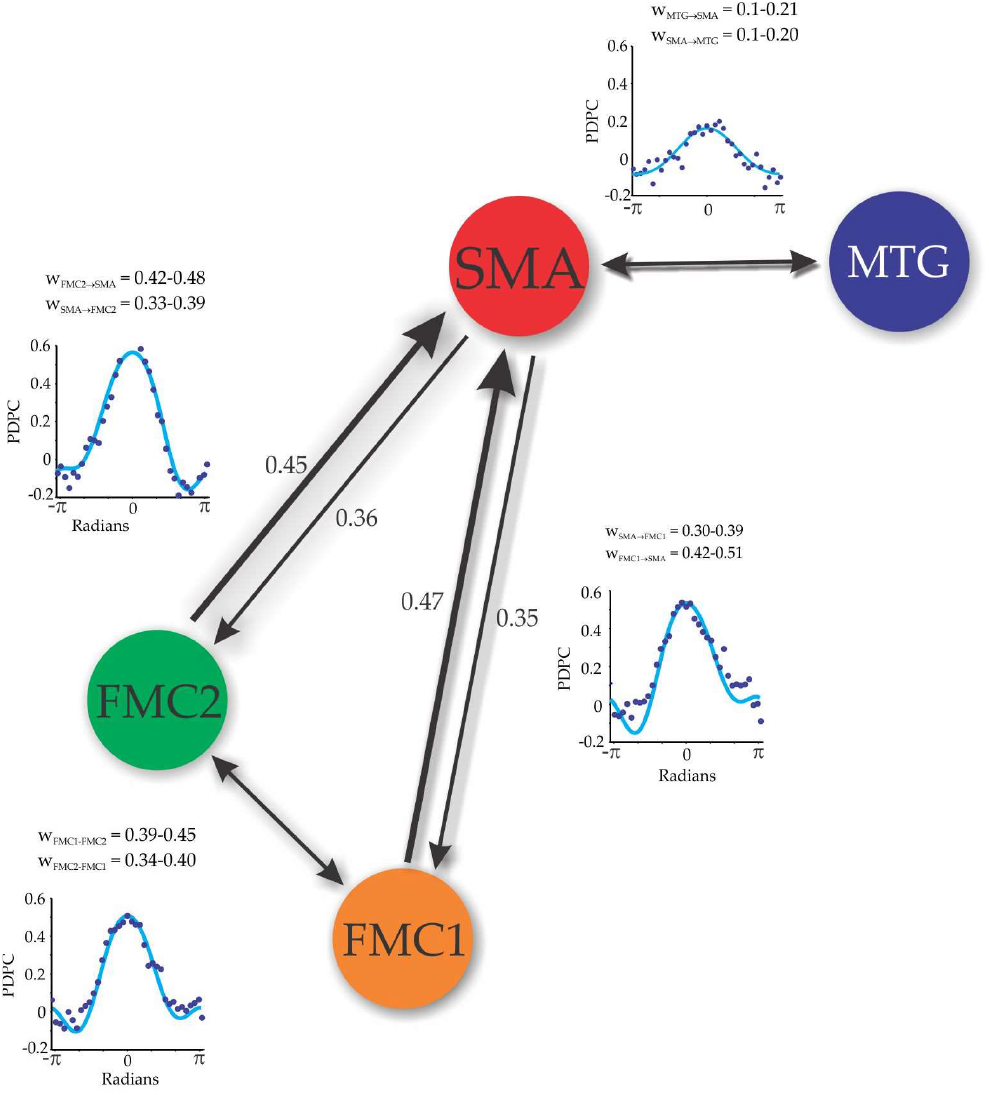
Fits of PDPC to bidirectional coupled oscillator model reveal stronger coupling from FMC to SMA. PDPC profiles for various electrode pairs (dark blue dots), and their associated fits (cyan) to a the bidirectional coupled model^30^ to obtain directional coupling weights (*w*). The weights are shown with their 95% confidence intervals. Directional arrows are shown with their associated weights (*w*) beside them. Where 95% confidence intervals overlapped, coupling was considered to be symmetrical.

Given the above interpretation of causal interactions at high-gamma, increased excitability (increased high-gamma power) in one region should more strongly drive excitability (high-gamma power) at a synaptically connected region. Indeed during language related tasks it has been demonstrated that electrodes displaying the greatest increases in high-gamma power are associated with the largest causal in-flow and outflows^13^. Interpreting these results on a finer temporal scale, one would expect to observe that high-gamma power and causal interactions are correlated on a moment to moment basis. To explore this potential relationship between power, and causal interactions for electrode sites that demonstration the largest high-gamma power increases, we uniquely correlated power fluctuations to causal interactions as a function of phase-relation. Such so called “phase-dependent power-causal correlations” were maximal within the γ_2_ range for both subjects (**Figure 6, and Supplementary Figure 10**), suggesting that increased high-gamma power at one site, has greater causal interactions at another, and this relationship is strongest near the good phase relationship between the sites.

**Figure 6:**
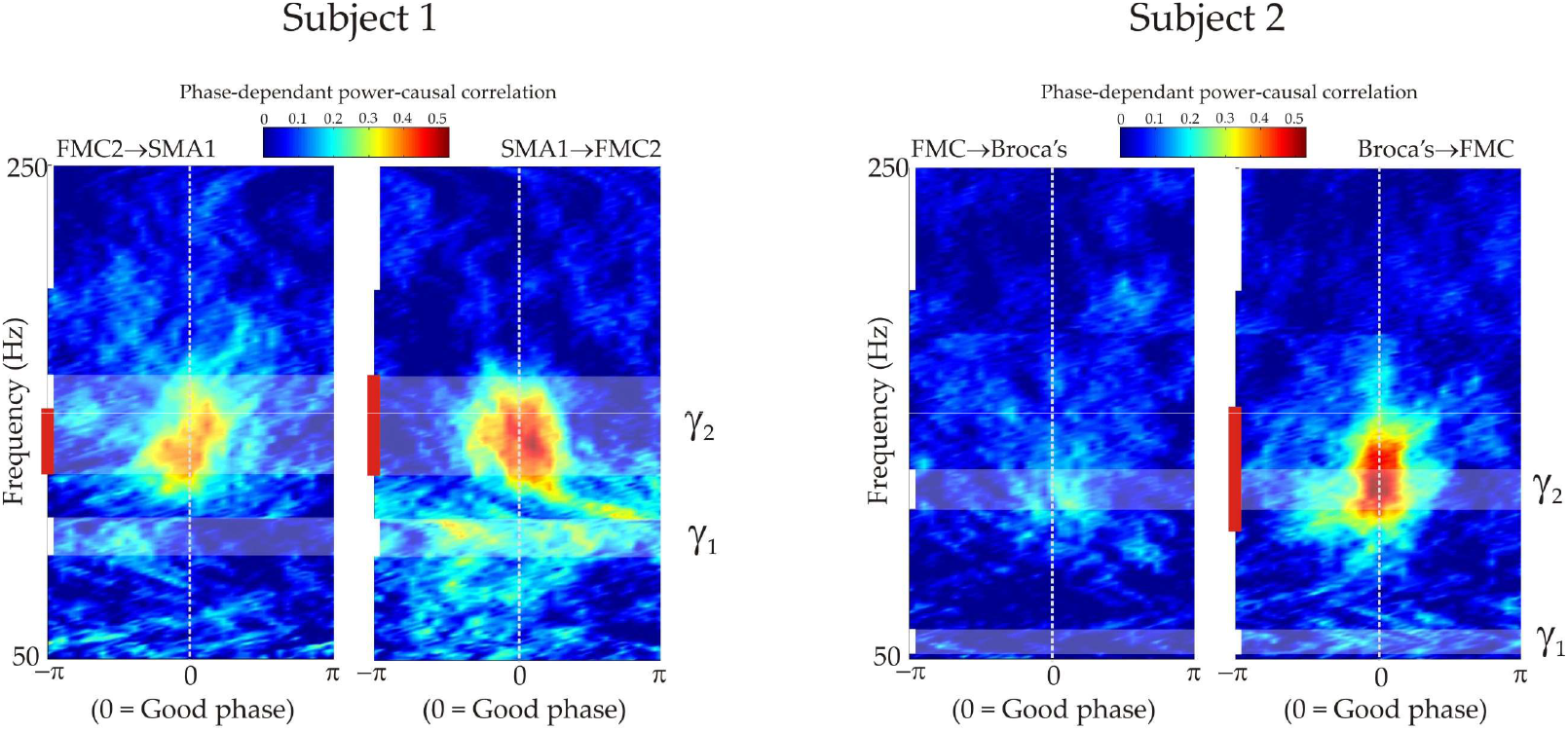
High-gamma phase, power and causal interactions are correlated. Analyses of electrode pairs that displayed the largest causal interactions for the two subjects. Phase-dependent power-causal correlations revealed significant correlations between causal interactions and power within the γ_2_ ranges that was strongest at the preferred phase relation between the two sites. Red bars represent frequency ranges of significance along the zero (dotted) line.

A physiological interpretation of causal interactions within the high-gamma frequency range might be that they represent a directed cascade of excitability through a network^13^. If indeed high-gamma is a proxy for temporal and spatial summation of spiking and synaptic activity, and this excitability (information) is transmitted axonally from on region to another, we would expect that the phase delays between regions are governed by axonal conduction velocity. The conduction velocities for long range connections (3 cm in distance) were 10 & 13 m/s for Subject 1, and for Subject 2, 5 m/s between Broca’s area and FMC (**Supplementary Figure 4**). These conduction velocities are well within those measured for myelinated fibers within the human CNS (7-24 m/s^46^), supporting the notion that cortico-cortical synchronization at high-gamma may represent “cascading”^13^ excitability from one node of the network to another.

## Discussion

In support of communication through coherence^3^ ESM identified regions that were both necessary for vocalization and which demonstrated the largest coherence between these sites. Surprisingly coherence and causal interaction were strongest at high-gamma frequencies, consistent with a previous report of causal interactions between ESM identified language sites within the high-gamma range^13^. Although typically communication through coherence is inferred by statistically significant increases in coherence during task related activity, our data here uniquely demonstrates that brain regions deemed necessary for vocalization, do indeed cohere. A number of observations suggest that our findings are not artifactual arising from volume or common reference problems, and are specific to the ESM positive sites. Firstly, we observed non-zero phase lagged synchronization that argues against our results being confounded by volume conduction, or common reference artifacts^19^. Secondly, language related tasks may be contaminated by temporalis muscle artifact, resulting in increased power in the high-gamma range. However, the spectral profile of such activity is typically broadband, with the greatest increases in the 20-80Hz range, and do not demonstrate peaks in the power spectrum^47^. The power spectra we observed during task related activity demonstrated clear peaks in ranges that are typically higher that the ranges over which muscle artifact generates power increases. Lastly, surrogate analysis utilizing contact pairs outside the SMA-FMC network demonstrated much lower, non-peaked coherence spectra, arguing for spatial specificity of our observation, and specifically that it was most profound within the electrodes that ESM identified as critical for vocalization.

Our findings are curious for two particular reasons. Firstly, unlike most studies, we observed peaks power spectra in the high-gamma range from iEEG recordings, which is not the norm. In this vain there is an extensive literature positing that the physiological basis of high-gamma activity is asynchronous spiking and post-synaptic potentials that when summed over space and time, generate increases in power that appear broadband band^11,48,49^. In this context it is extremely important to note that in this study we present data of only two patients. When performing such experiments in other patients, we like others have observed non-peaked power spectra in the majority of such patients.

We have previously shown that supragranular layers of human neocortex, the source of long range cortico-cortical projects^50^ can generate high-gamma activity^51^ suggesting that neuronal activity in supragranular layers may start the process anew at distant sites^2^. The origin of high-gamma activity from such feed-forward laminae of the cortex suggests that communication through coherence at high-gamma frequencies may not be symmetrical. The phase delays and aDTF estimate of causal interaction support this conclusion, and suggest that high-gamma activity establishes a temporary standing directed wave^39^, that is phase delayed by axonal conduction times. Furthermore, phase-dependent power-causal correlations suggest that temporal precision, power, and causal interaction are intimately linked to one another within the high-gamma range (**Figure 5**). On the whole, high-gamma and its amplitude modulation may not only index local cortical processing^10,21,29,34,48,52–56^ but as well long range communication, it’s direction, and causal interactions at least within language processing areas^13,14^, and possibly throughout the brain.

In summary our findings demonstrate that under some specific circumstances long-range directed causal influences may at times be mediated by phase-coherence high-gamma oscillations. Together with similar observations within language networks^13,14^, and a paucity of such findings outside such regions^12^, our data suggests that phase-coherent high-gamma activity may be somewhat specific to speech production and language areas of the human brain.

### Conclusions

Power increases in the high-gamma range may in some situations imply local synchronous population activity, large-scale cortico-cortical interactions^13,14,57^, and participation in a common and temporally unfolding cognitive activity within networks of the human brain particularly those related to language.

## Author Contributions

TAV performed the surgeries, designed the experiments, conducted the stimulation mapping, wrote the analysis software, analyzed the data, and wrote the manuscript. BG wrote analysis software, and edited the manuscript.

## Acknowledgements

We would like to thank T. Womelsdorf and K. Hoffman for very helpful comments regarding the manuscript. TAV would like to acknowledge the mentorship of G.A. Ojemann who taught him the technique of stimulation mapping utilized in this study.

The author declare no competing financial interests.

