## Supplementary material and figures for "Long range, high-gamma phase coherence in the human brain during overt and covert speech"

### Supplementary Materials:

#### Subject information and task details:

**Subject 1** was a 22 year old male, with medically intractable epilepsy for 5 years. His 3T MRI was normal, and he underwent resection of the left frontal pole. His irritative zone ictal onset zones were confined to the frontal polar region. Pathology revealed gliosis. Neither interictal activity nor ictal onsets were recorded from any of the contacts that were used for analysis. Temporal contacts were not mapped.
Within the inter-hemispheric fissure stimulation at the "leg" electrodes with 4.5 mA resulted in hip and knee flexion although 3 mA induced subjective leg numbness , whereas at the "arm" electrodes a combination of arm and leg flexion occurred, albeit predominantly arm. Within the "speech arrest" electrodes the posterior pair obtained speech arrest at 8.5 mA while the anterior two contacts arrest was obtained at 3.5 mA. Anatomically the most posterior inter-hemispheric contact was in line with the central sulcus placing it on the leg area of the primary motor cortex, anterior to primary sensory area of the leg. More anterior stimulation resulted in arm movement and speech arrest attesting to the location of these electrodes with the SMA (Fried et al. 1991). FMC1 electrodes required less current (5 mA) than FMC2 electrodes (12 mA) to elicit speech arrest. Although the more anterior electrodes may have rested on pre-motor cortices, for simplification all electrodes within this region will be referred to as face motor cortex (FMC). No vocalization were obtained during mapping, and all speech arrest sites resulted in the same objective observation of speech arrest.

**Subject 2** was a 59 year old male, with medically intractable epilepsy for 20 years. Left frontal pole ulegyria was apparent on 3T MRI. His ictal onset zones was contained within the left frontal lesion, however irritative zones included the lesion and a rim of surrounding tissue, orbito-frontal cortex, and the left anterior temporal and right posterior temporal regions. He underwent resection of the left frontal pole lesion and surrounding rim of tissue, with pathological analysis revealing ulegyria and gliosis. Mapping proceeded as with subject 1, however over the left frontal convexity within a 2x8 cm a variety of speech alterations including dysarthria, decreased fluency, and speech arrest were obtained (see **Figure S4**).

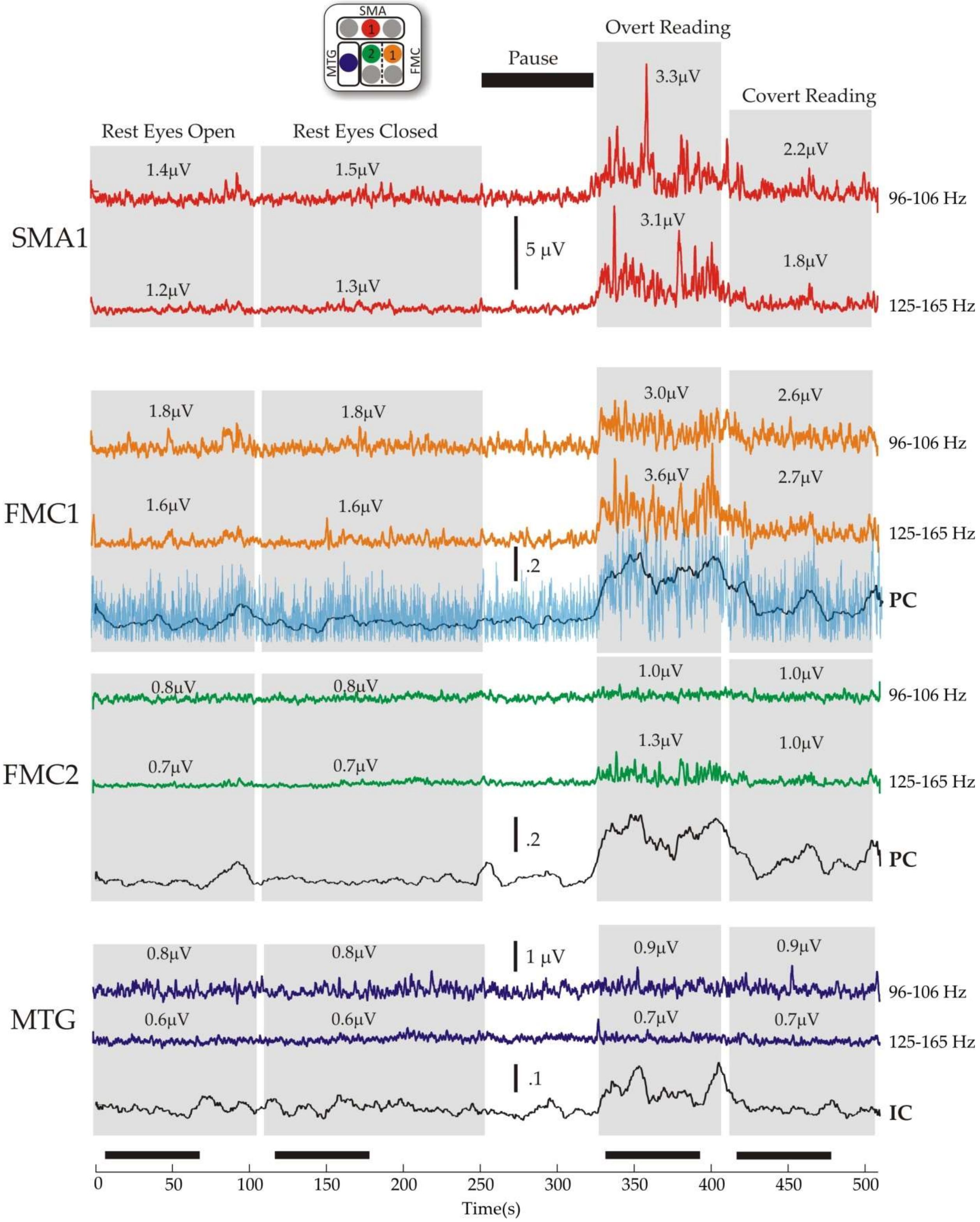
SMA
MTG
FMC
Pause
Overt Reading
Covert Reading
Rest Eyes Open
Rest Eyes Closed
SMA1
FMC1
FMC2
MTG
96-106 Hz
125-165 Hz
PC
IC
5 μV
1 μV
Time(s)

**Figure S1| Experimental time course, power changes, and synchronization  for subject 1.** Grey areas represent various experimental time periods (labeled above) over which behavioral tasks were performed, and black bars at the bottom of the figure represent the 60s regions over which data was analyzed for each condition.  The color of the traces correspond to the legend at the top, and the electrode colors to the that in **Figure 1**. Two amplitude traces are displayed for four different electrodes by taking the absolute value of the Hilbert transformed band pass filtered trace, and heavily smoothing for display using a sliding 800ms window.  The frequency ranges for the band pass filtering was obtained from the power spectra for the SMA1 electrode (**see Figure S2**).  The numbers above the traces refer to the mean amplitude over the 60s time period. Below the amplitude time courses are show synchronization time courses, for FMC1, FMC2, and MTG to the SMA1 electrode.  For FMC1 and FMC2 PC is displayed, whereas for MTG IC is displayed, as phase-coherence did not identify significant synchronization between these two electrodes (**see Figure 4**). The light blue trace superimposed on the PC time course for FMC1 represents the unsmoothed PC time series, highlighting the non-stationary, albeit sustained increases in synchronization.

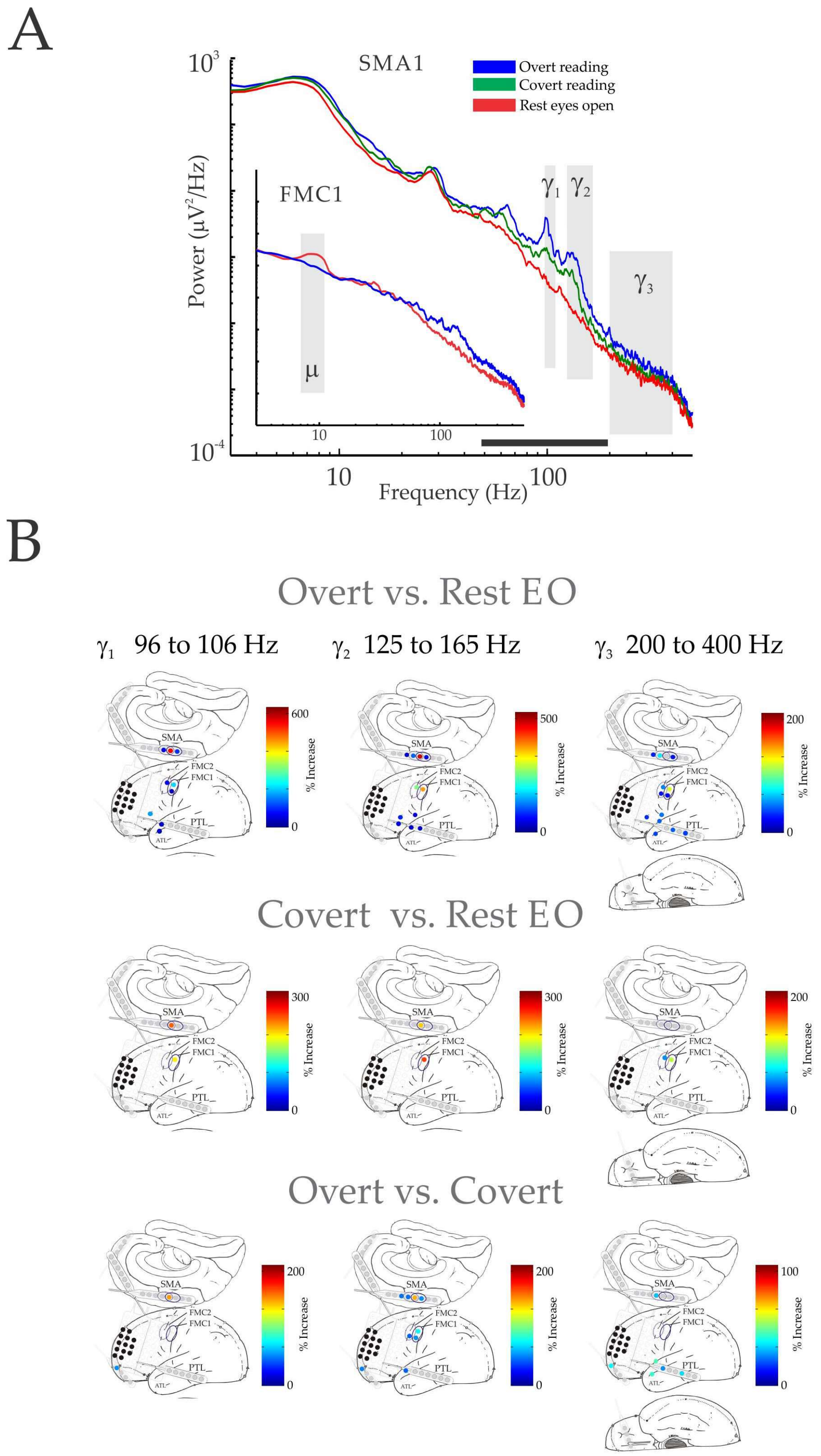
A
SMA1
Overt reading
Covert reading
Rest eyes open
Power (μV²/Hz)
Frequency (Hz)
FMC1
μ
γ1
γ2
γ3
B
Overt vs. Rest EO
γ1 96 to 106 Hz
γ2 125 to 165 Hz
γ3 200 to 400 Hz
Covert vs. Rest EO
Overt vs. Covert
SMA
FMC2
FMC1
PTL
ATL
% Increase

**Figure S2 | Identification of gamma frequency ranges, and summary of power changes during reading.** **(A)** Power spectra for the SMA1 and FMC1 (inset) electrodes. $\gamma_1$(96-106Hz), and $\gamma_2$ (125-165Hz) (grey bars) were defined by regions of significant power increases between overt as compared to covert reading, while $\gamma_3$ (200– 400Hz) was arbitrarily set since this range has been shown to carry unique information and likely to arises from different neuronal mechanisms (Jones et al. 2000; Baker et al. 2003; Zhuang et al. 2010). The power spectrum for FMC1 displays a typical peak in μ range, that dissipated during overt reading consistent with its location over the motor strip. The black bar represents the region of significant power increase for overt reading as compared to the rest eyes open state. (**B)** Summary of significant power increases for the various gamma ranges, for all electrodes from which recordings were obtained. Only the posterior half of the grid was recorded from (separated from the anterior portion by a vertical line). The black electrodes represent the epileptogenic region – as resection of this region resulted in seizure freedom. The color bar corresponds to percent increase from the second condition of the comparison. As three different comparisons were performed alpha was set to 0.017 for FDR correction. The FMC1, FMC2, and SMA1 electrodes are labeled. The ovals correspond to the combination of mapped electrode pairs within the SMA-FMC network (**Figure 1**).

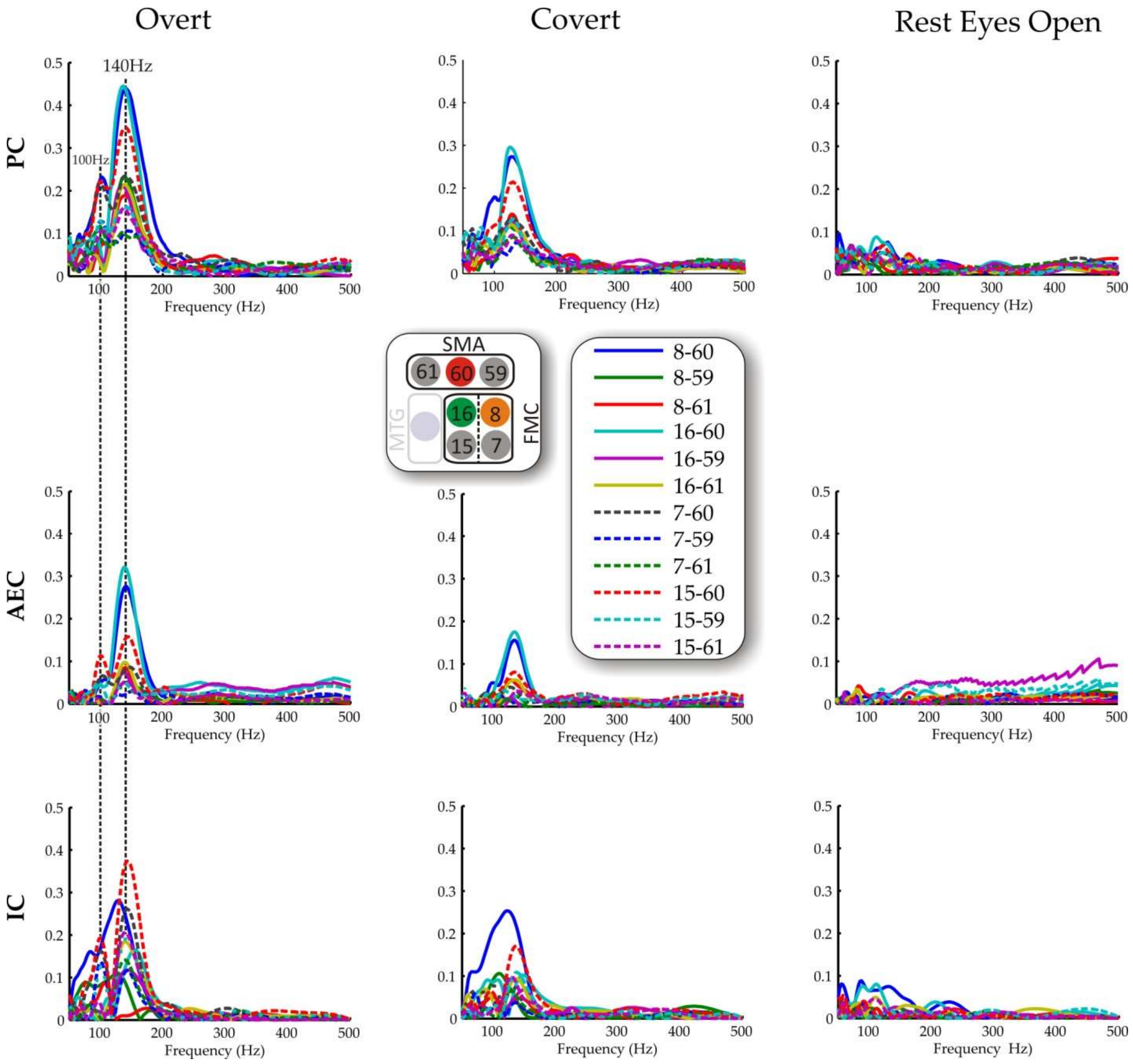


**Figure S3 | Individual entrainment profiles for the twelve inter-areal pairs arising from the SMA, and FMC**. Profiles are shown for the three different entrainment measurements. Contact 8 is FMC1, contact 16 is FMC2, and contact 60 is SMA. MTG is not included in these profiles.

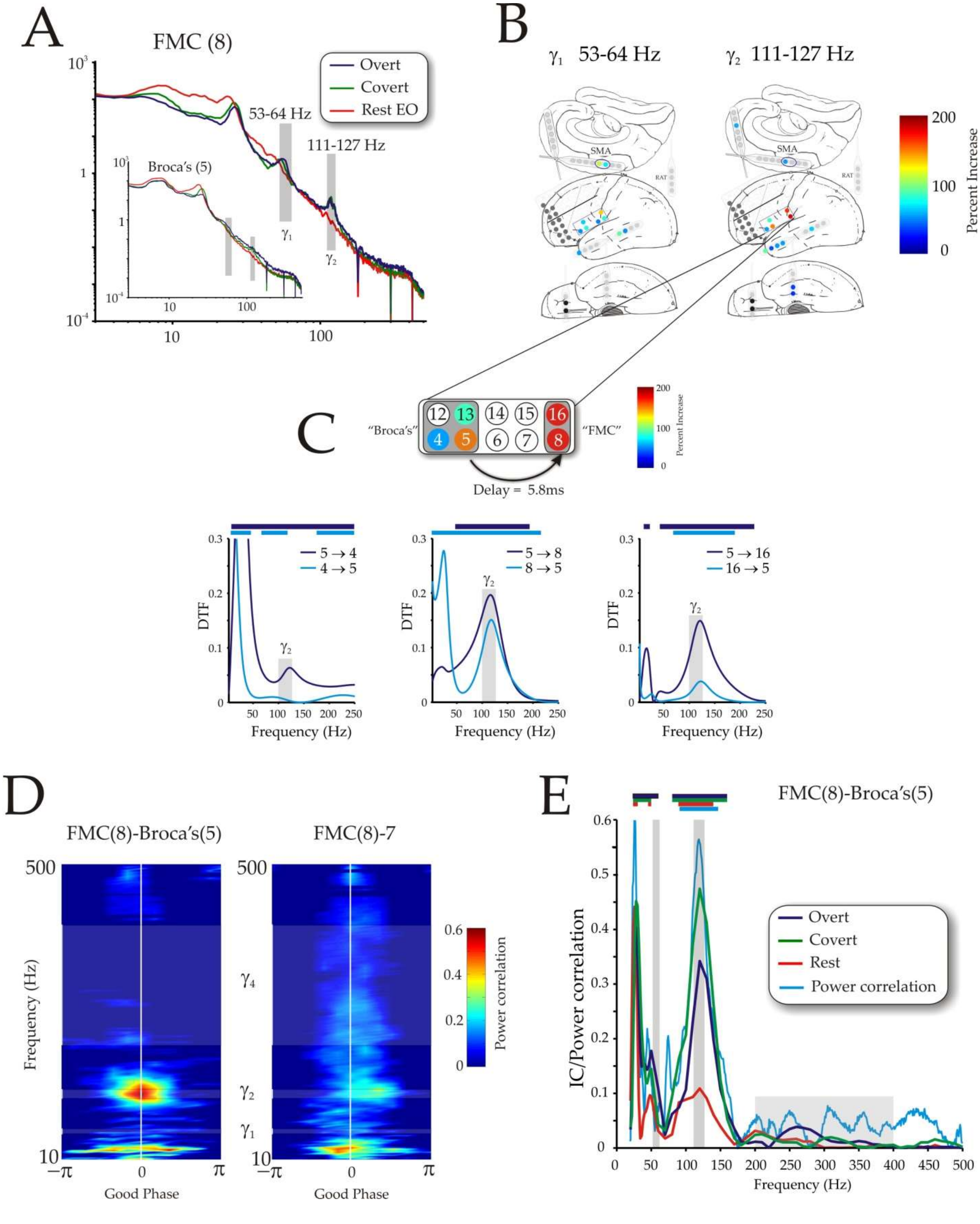
A
FMC (8)
Overt
Covert
Rest EO
53-64 Hz
111-127 Hz
Broca's (5)
B
γ1 53-64 Hz
γ2 111-127 Hz
SMA
Percent Increase
C
"Broca's"
"FMC"
Delay = 5.8ms
DTF
Frequency (Hz)
5 → 4
4 → 5
5 → 8
8 → 5
5 → 16
16 → 5
D
FMC(8)-Broca's(5)
FMC(8)-7
Frequency (Hz)
Good Phase
Power correlation
E
FMC(8)-Broca's(5)
IC/Power correlation
Overt
Covert
Rest
Power correlation
Frequency (Hz)

**Figure S4 | Subject 2 summary.** **(A)** Power spectra for electrode within FMC (electrode 8) which when paired with electrode 16 during stimulation mapping obtained speech arrest. Inset displays power spectra for electrode within Broca's area (electrode 5) which when paired with electrode 13 during stimulation mapping obtained dysarthria. $\gamma_1$ (53-64Hz), and $\gamma_2$ (111-127Hz) bands were defined from the FMC electrode when comparing covert reading to rest. No significant power changes were noted between overt and covert reading. Notch filtering was performed at 180, 300, and 480Hz (this was not required in subject 1). Note the clear peaks, in the gamma frequency range. (**B)** Summary of power changes comparing covert reading to the rest state within the identified frequency bands for all recorded electrodes. For the grid, recordings were obtained from the 32 electrodes below the black line. The gray electrodes represent the ictal onset zone. Color bar represents the percentage increase above the rest state. (**C)** Expanded view of the analysis region. The grayed regions represent areas where speech disturbances were obtained. In Broca's area dysarthria was obtained without speech arrest, whereas within the FMC speech arrest occurred. The delay of 5.8 ms between the indicated electrodes reflected a conduction velocity of 5 m/s that was computed for the $\gamma_2$ frequency range, using a linear distance of 3cm. This delay is consistent with cortico-cortical evoke potentials between Broca's and motor cortex that peak between 2.6-6ms (*32*). Directed transfer function analyses are presented below. Grey bars identify the $\gamma_2$ frequency range. (**D)** Phase dependent power correlations for indicated electrode pairs during covert signing. Note the weaker correlations within the $\gamma_2$ range for immediately adjacent electrodes. (**E)** Frequency profiles for IC and phase-dependent power correlations between FMC and Broca's area. The colored bars indicate regions of significance above chance for the various measures and conditions. The grey bars correspond to the two defined gamma ranges.

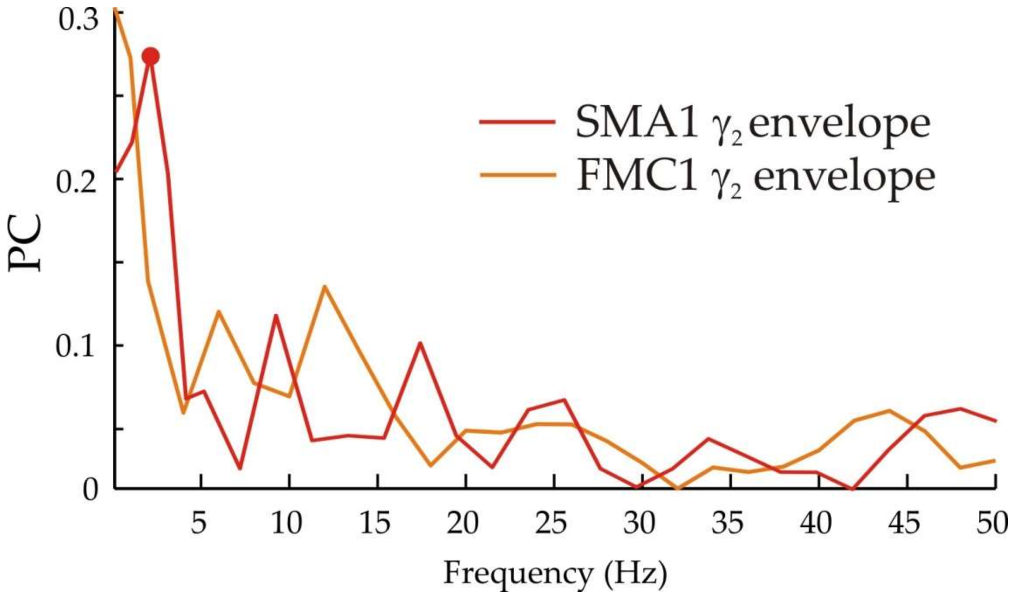


**Figure S5 | $\gamma_1$ and $\gamma_2$ synchronization to LFPs during overt reading.** Envelopes of the $\gamma_1$ and $\gamma_2$ oscillations were obtained from the Hilbert transformed band pass filtered time series of FMC1 and SMA during overt reading. The PC between these time series and local field potential were computed for frequencies between 2 and 50Hz. Symbol represents significant increase from chance using bootstrap techniques. Significance was not obtained for the other behavioral conditions.

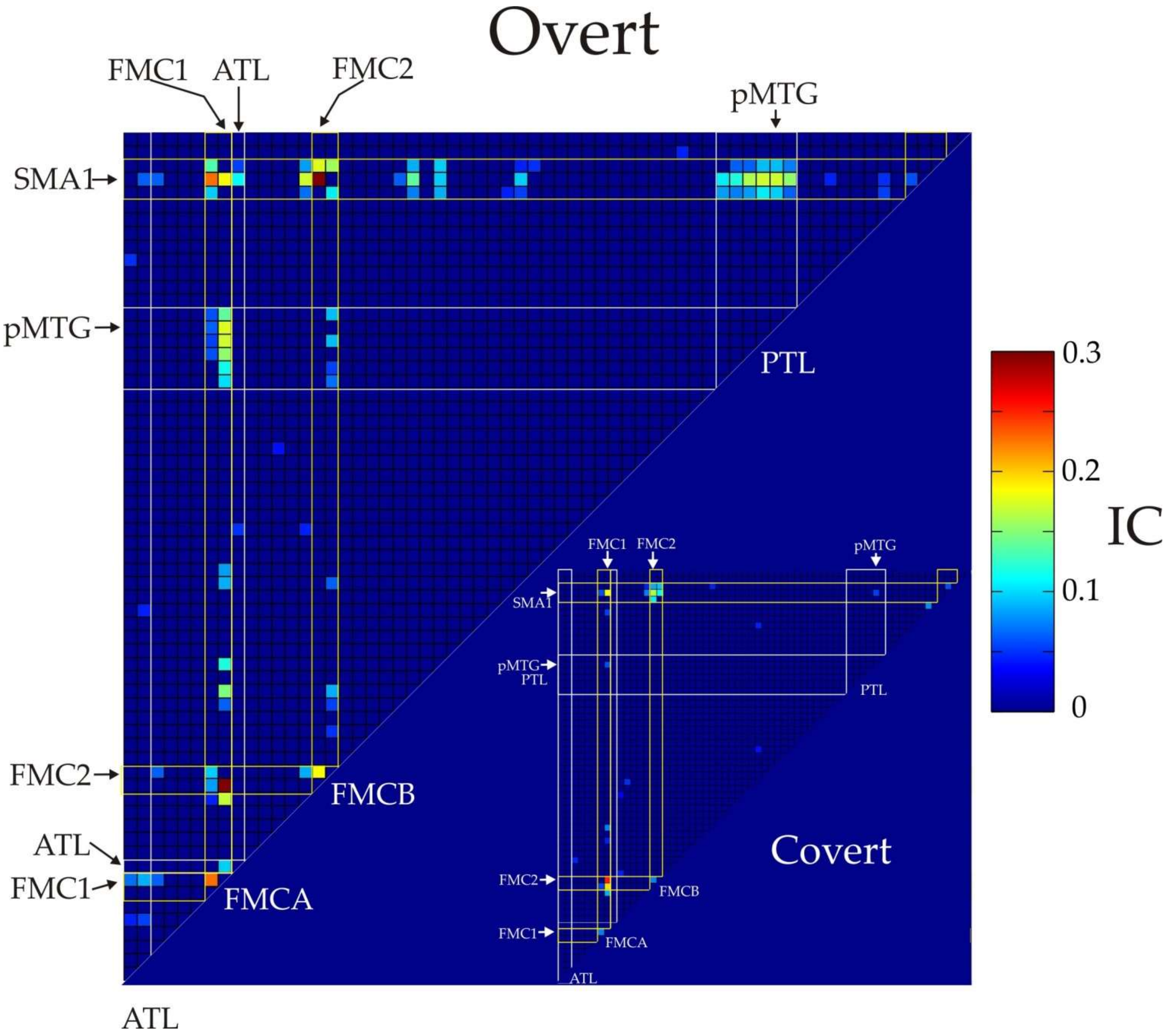


**Figure S6 | IC adjacency matrices for the $\gamma_2$ frequency band during overt and covert reading.** The various contact groups are labeled and identified by white or yellow (SMA and FMC regions) boxes. FMCA and FMCB refer to the contact pairs 7&8, and 15&16 respectively (**see Figure 1 and S3**). Individual electrodes are identified with an arrow and associated text. As can be appreciated from this matrix the majority of electrodes are connected directly to the SMA-FMC network. Only 8 electrodes are not directly connected to the SMA-FMC network, although 6 of these 8 are indirectly connected via an intervening electrode.

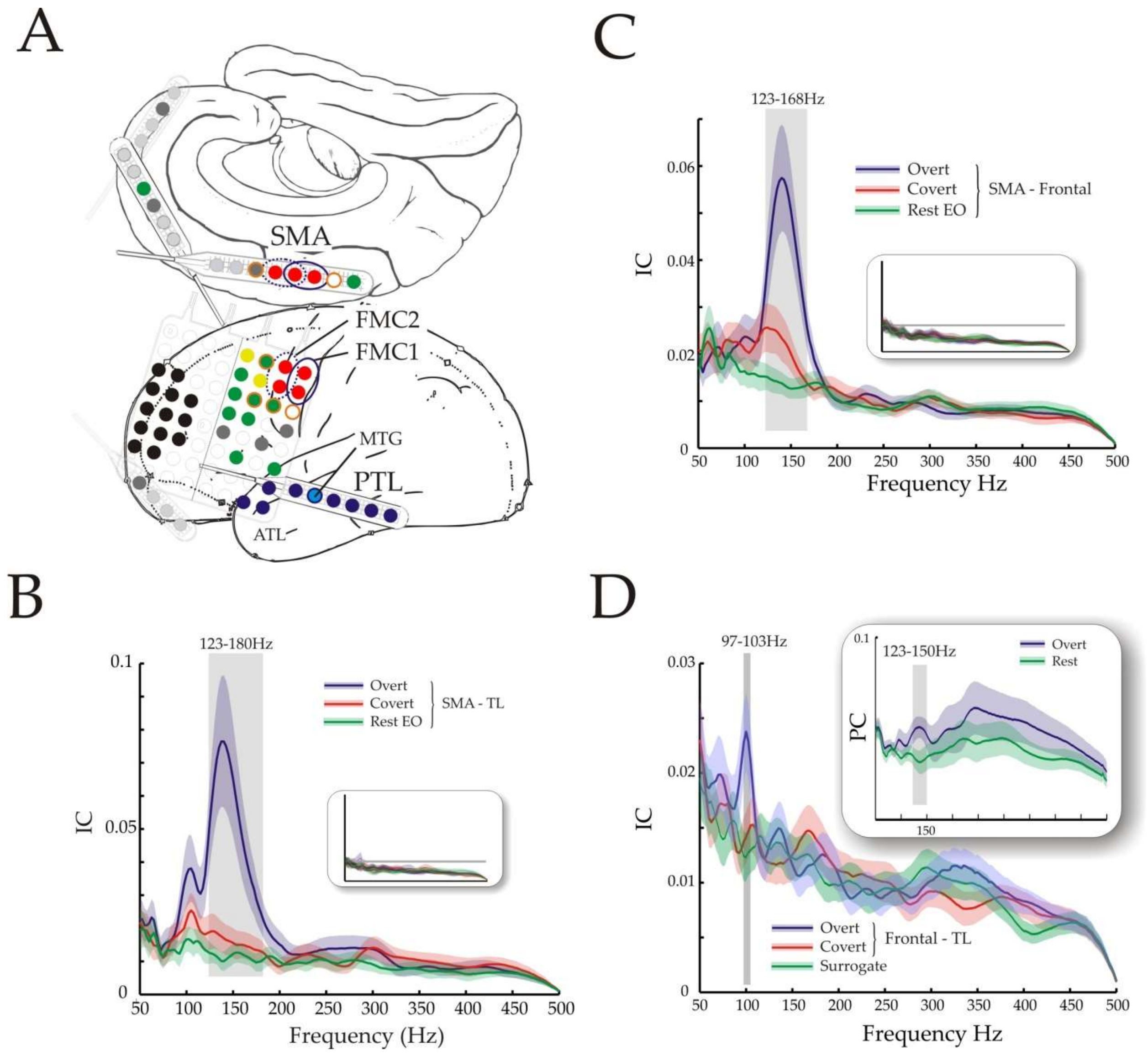


**Figure S7 | Sub-network analysis of $\gamma_2$ reading network.** (**A**) Schematic of electrodes entrained within the $\gamma_2$ frequency "network" during overt reading. Of those electrodes entrained directly to the SMA-FMC network (red electrodes), blue electrodes are within the temporal lobe (TL: which includes the ATL and PTL electrodes), and green electrodes are distributed throughout the frontal lobe. The MTG electrode is depicted with a light blue boarder and was the only TL electrode that displayed entrainment to the SMA-FMC network during both overt and covert reading (**Figure 4**). Those electrodes with an orange border were at the periphery of the SMA-FMC network and used for the insets of B & C. Grey electrodes are those not directly connected to the SMA-FMC network (8 in total). One grey electrode in the orbital frontal cortex is not shown for clarity. Of the grey electrodes three were neither directly nor indirectly connected to the SMA-FMC network (one grey orbitofrontal contact is not displayed). Black electrodes ictal onset zone – recordings were not obtained from these electrodes. Lastly, yellow electrodes were contaminated by noise and not included for analyses. (**B**) Frequency dependent synchronization between FMC and the TL. The $\gamma_2$ synchronization region dissipates during covert reading, in contrast to the SMA-FMC (**Figure 1D**). Inset displays frequency dependent synchronization for electrodes at the periphery of the SMA-FMC network (electrodes with orange boarder) with TL electrodes. Superimposed are the time-shifted surrogate frequency dependent synchronization profiles. Horizontal grey line depicts an IC of 0.02 for reference.

(**C)** Frequency dependent synchronization for pairs arising between all SMA-FMC network and frontal lobe electrodes. Inset is same analysis but for electrodes peripheral to the DSN with frontal electrodes and surrogates. Horizontal grey line depicts an IC of 0.02 for reference. **(D)** Frequency dependent synchronization profile between the frontal lobe and temporal lobe. Regions of significance are demarcated by grey bars.

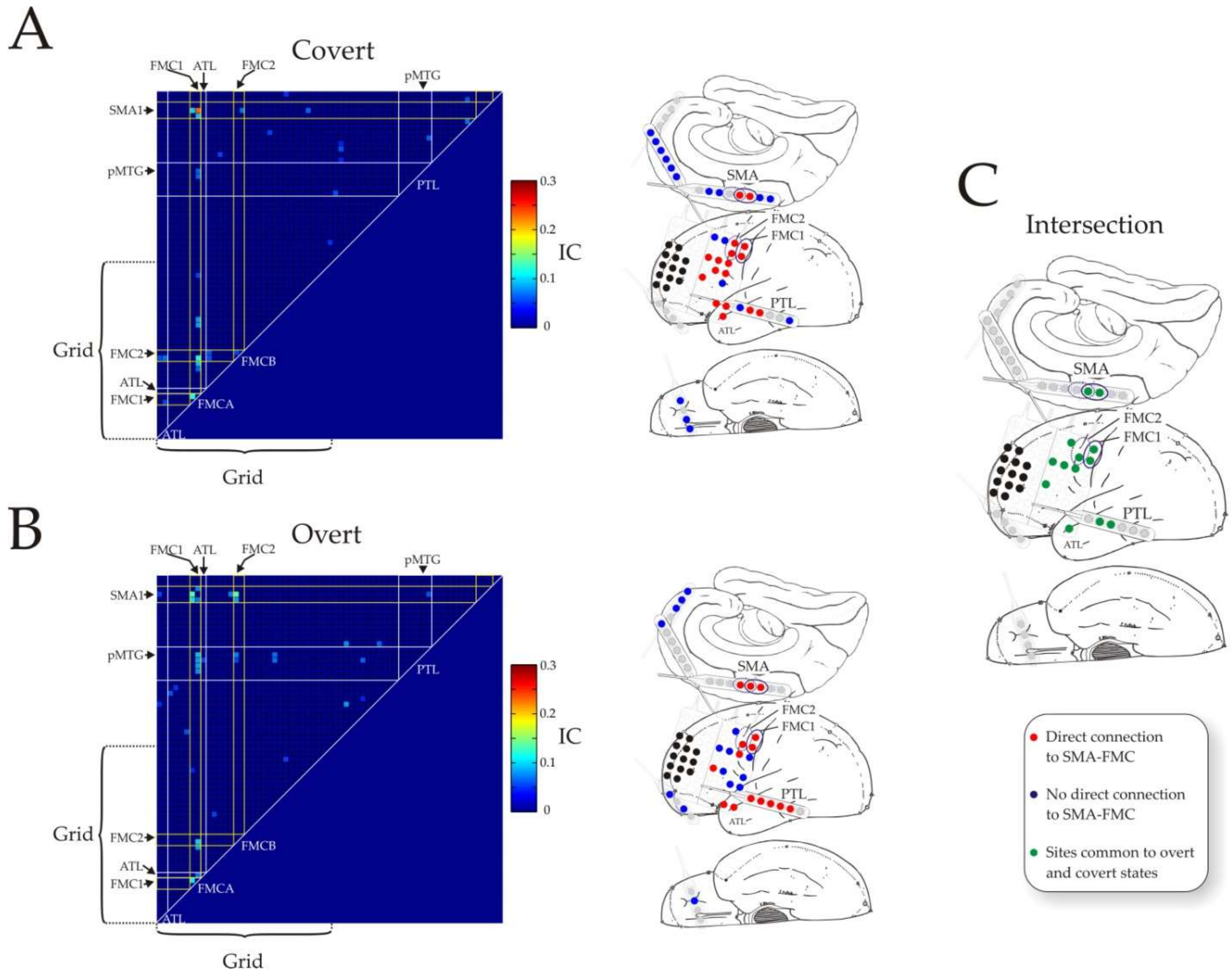


**Figure S8 | $\gamma_1$ reading network:** Network was constructed using band pass filtered signals between 96-106 Hz in a fashion analogous to that used for the $\gamma_2$ network (**Figure S7**). **(A)** During overt reading. (B) During covert reading. (C) The intersection of the networks in (A) and (B). Of particular note is the continued inclusion of the middle temporal gyrus that is common to both overt and covert reading, as well as prefrontal regions that are entrained within the $\gamma_2$ network as well.

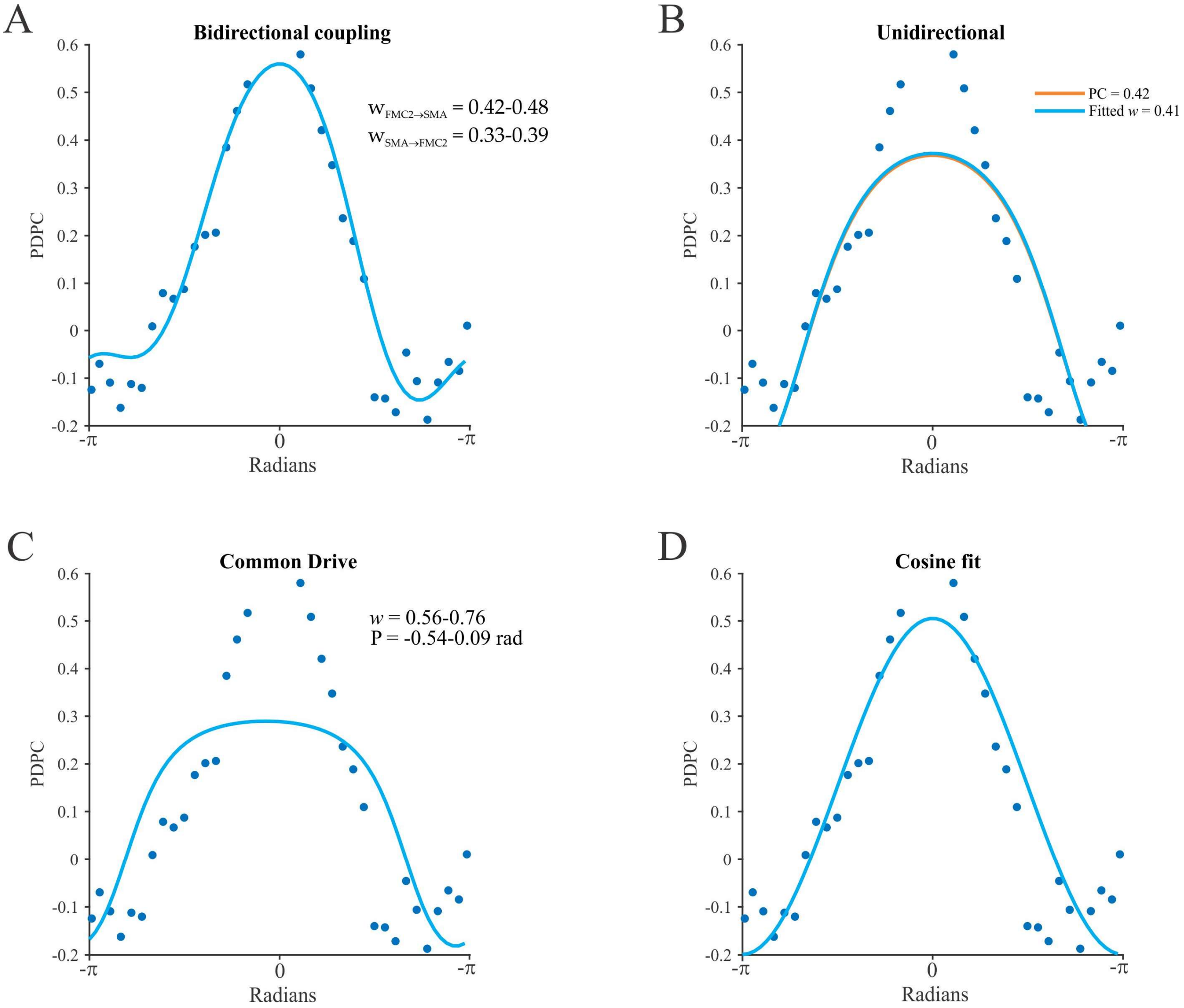


**Figure S9 | Model fits to FMC2 and SMA electrode phase-dependent power correlations.** Model parameters estimated using closed form expressions for PDPC. A) Bi-directional model. The coupling strengths are denoted by *w*. B) Unidirectional fit (blue), and *w* estimated from phase coherence (PC). C) Common drive model where ϕ is the coupled phase difference between oscillators. D) Cosine fit used to in original description of PDPC (Womelsdorf et al. 2007).

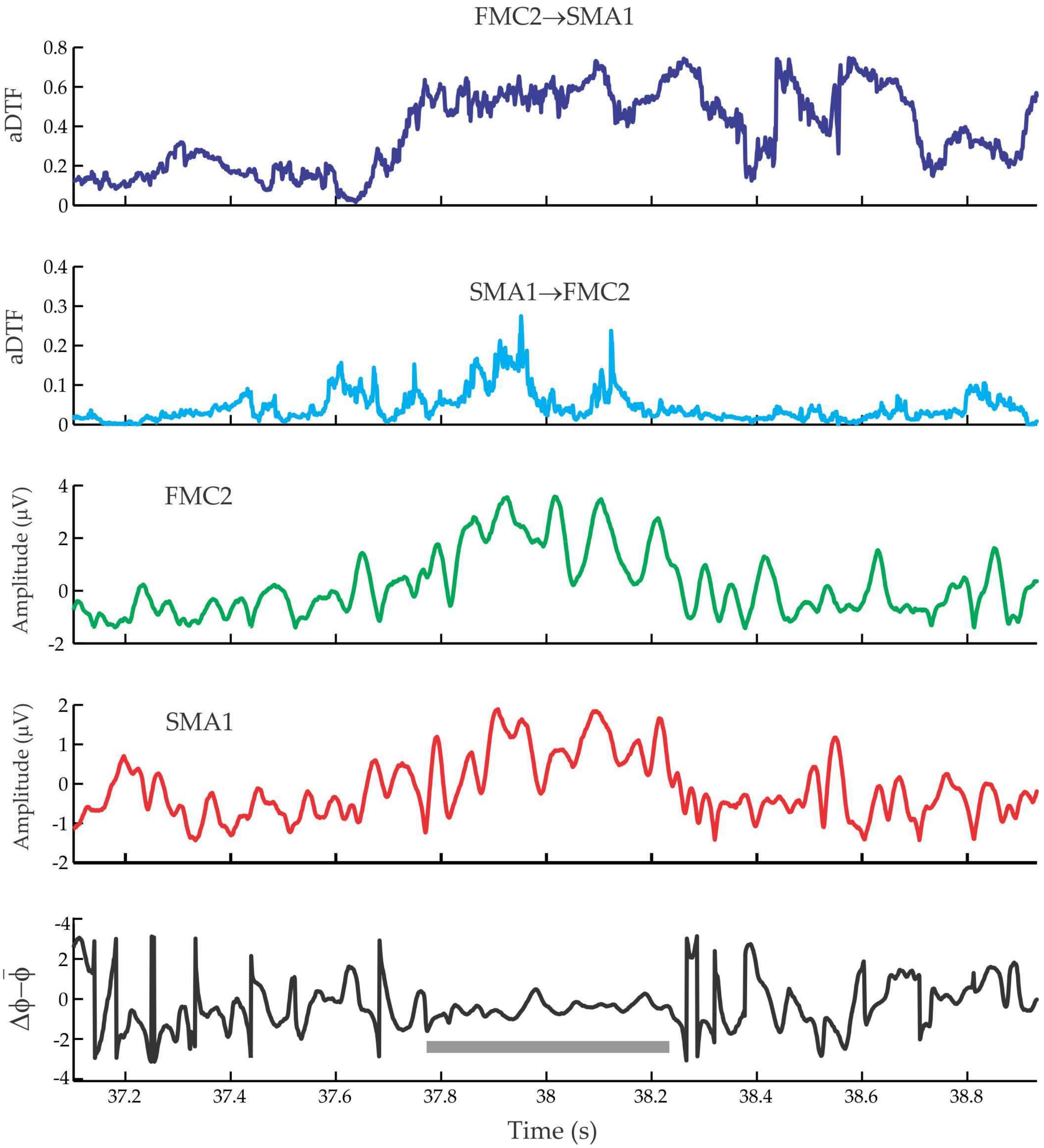


**Figure S10 | Time course for power, causal interactions, and phase-relations used to compute phase-dependant power-causal correlations.** From subject 1 FMC2 and SMA1 time series of the various measures at 140Hz are used as an exemplar. These short time segments exemplify that when the phase-relation hovers near zero there is an increase in power and causal interactions.

Baker SN, Curio G, Lemon RN. 2003. EEG oscillations at 600 Hz are macroscopic markers for cortical spike bursts. J Physiol. 550:529–534.
Fried I, Katz A, McCarthy G, Sass KJ, Williamson P, Spencer SS, Spencer DD. 1991. Functional organization of human supplementary motor cortex studied by electrical stimulation. J Neurosci. 11:3656–3666.
Jones MS, Jones MS, MacDonald KD, MacDonald KD, Choi B, Choi B, Dudek FE, Dudek FE, Barth DS, Barth DS. 2000. Intracellular correlates of fast (>200 Hz) electrical oscillations in rat somatosensory cortex. J Neurophysiol. 84:1505–1518.
Womelsdorf T, Schoffelen JM, Oostenveld R, Singer W, Desimone R, Engel AK, Fries P. 2007. Modulation of neuronal interactions through neuronal synchronization. Science (80- ). 316:1609–1612.
Zhuang J, Truccolo W, Vargas-Irwin C, Donoghue JP. 2010. Decoding 3-D reach and grasp kinematics from high-frequency local field potentials in primate primary motor cortex. IEEE Trans Biomed Eng. 57:1774–1784.
